# A genetic locus in the gut microbe *Bacteroides thetaiotaomicron* encodes activities consistent with mucin-O-glycoprotein processing and plays a critical role in *N*-acetylgalactosamine metabolism

**DOI:** 10.1101/2024.02.01.578401

**Authors:** Didier A. Ndeh, Sirintra Nakjang, Kurt J. Kwiatkowski, Nicole M. Koropatkin, Robert P. Hirt, David N. Bolam

**Affiliations:** Division of Plant Sciences, School of Life Sciences, University of Dundee, Dundee, United Kingdom; Precision Medicine Centre of Excellence, Queen’s University, Belfast, Belfast, United Kingdom; Department of Microbiology and Immunology, University of Michigan Medical School, Ann Arbor, MI, USA; Biosciences Institute, Medical School, Newcastle University, Newcastle upon Tyne, NE2 4HH, UK

## Abstract

It is increasingly appreciated that members of the gut microbiota are key modulators of human health and the status of major diseases including cancer, diabetes and inflammatory bowel disease. Central to their survival is the ability to metabolise complex dietary and host-derived glycans including intestinal mucins. The latter are critical components of the gut epithelium glycocalyx and mucus barriers, essential for microbiota-gut homeostasis and protection from infections by pathogens. The prominent and model human gut microbe *Bacteroides thetaiotaomicron (B. theta)* is a versatile and highly efficient complex glycan degrader thanks to the expansion of gene clusters termed polysaccharide utilisation loci (PULs) in its genome. While the mechanisms for several singular dietary glycan-induced PULs have been elucidated, studies on the 16-18 mucin-induced PULs in *B. theta* significantly lag behind. A combination of the scale and complexity of *B. theta* transcriptomic response to mucins and complex glycan configurations of mucins represent major hurdles for the functional characterisation of the mucin induced PULs. As a result, there is very limited knowledge on how mucin metabolism is coordinated in *B. theta* and what specific PULs, genes and metabolites are critical for mucin-*B. theta,* and more generally mucin-microbiota interactions and their importance in microbiota-gut homeostasis. Here we show that a mucin inducible PUL BT4240-50, (i) encodes activities consistent with a machinery that couples the processing of mucin-O glycan glycoproteins with the metabolism of *N*-acetylgalactosamine (GalNAc), an abundant mucin O-glycan sugar; (ii) is important for competitive growth on mucins *in-vitro*; (iii) encodes a key kinase enzyme (BT4240) that is critical for GalNAc metabolism and (iv) has related PULs encoded by a range of prominent *Bacteroides* species in the human gut. Furthermore, BT4240 kinase was also critical for glycosaminoglycan metabolism, thus extending the PULs function beyond mucins. Our work advances our understanding of the vital metabolic processes that govern mucosal glycoprotein metabolism and by implication, a key aspect of host-microbiota interactions at mucosal surfaces and highlight GalNAc as a key metabolite targeted for competitive growth.

## Introduction

The majority of the human gut microbiota resides in the colon, with the colonic microbiota increasingly recognized as a metabolic organ that profoundly influences numerous aspects of human biology [1]. Critical to microbiota-colon homeostasis is the mucus layer that provides protection for the underlying epithelia from the huge microbial load [2–4]. Colonic mucus is composed of two discrete layers, the looser outer layer is a niche for some members of the microbiota and is associated with the essentially microbe-free and more rigid and impenetrable inner layer [2–5] (Fig.1A). These two strata are essential in maintaining digestive and immunological homeostasis where human enterocytes and immunocytes coordinate their activities with resident microbes for mutually beneficial outcomes [4, 6, 7].

**Figure 1:**
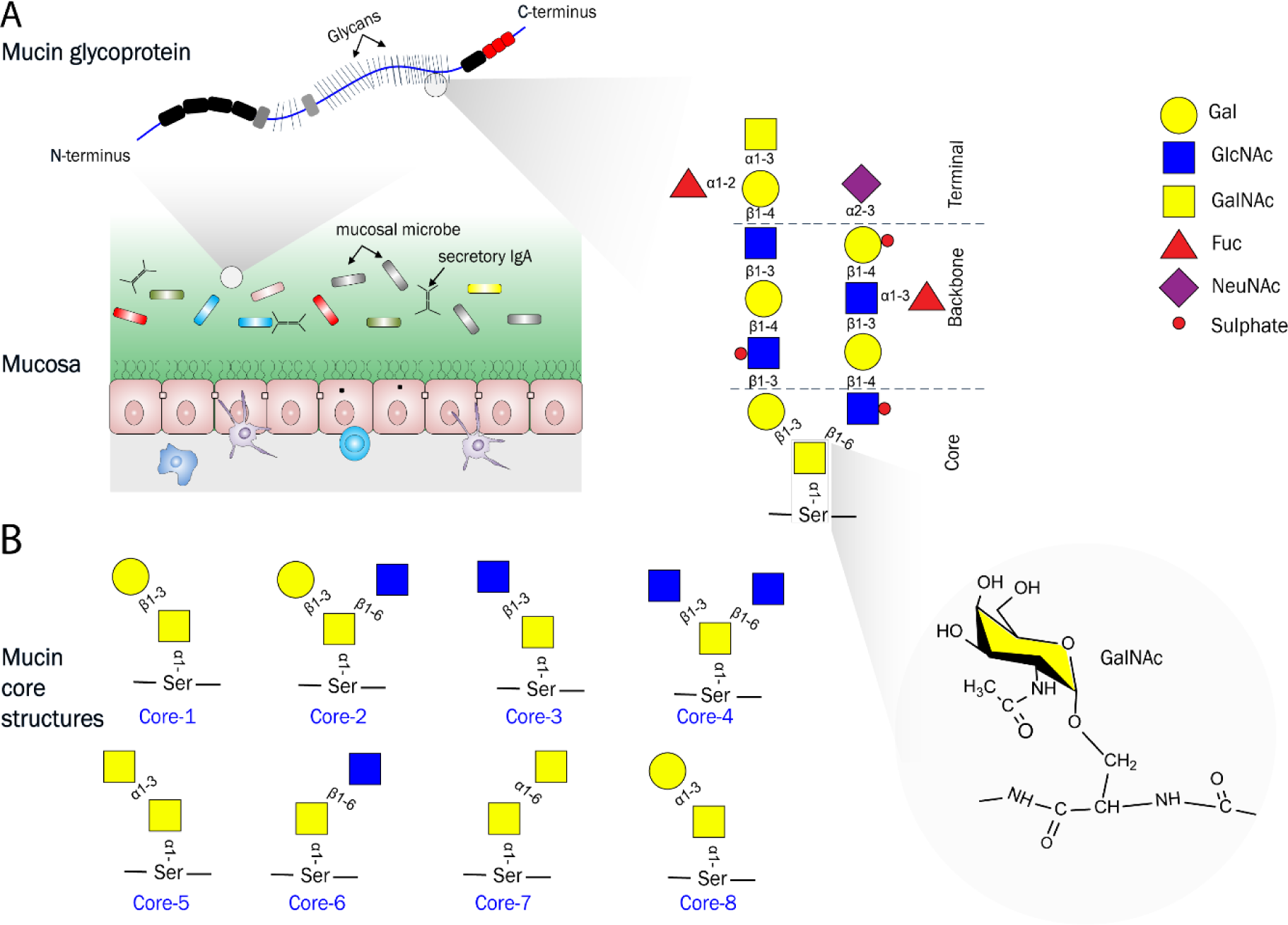
Mucins and various mucin cores structures showing the context and location of the core and terminal sugar N-acetylgalactosamine (GalNAc) **A:** Mucins are the major organic components of mucus on mucosal surfaces and consist of a protein backbone heavily decorated by O-linked glycans resulting in a ‘bottle brush’-like appearance [2, 10, 11, 40]. GalNAc is the hallmark and core of mucin-O type sugars forming the first O-glycosidic linkage to the peptide with the side chain of serine or threonine. GalNAc is also present as a terminal decoration of mucin side chains such as Blood group A. **B:** O-glycan side-chains are highly variable, but are composed of eight known core structures[12], with cores 1-4 most common in intestinal mucins. Core-3 is common in colonic mucin (mainly MUC2), whereas core-1 is more common in the secreted mucins of the upper digestive tract (e.g. MUC5AC, MUC6) [11, 40, 58].

Complex glycans are the major nutrient source available to the colonic microbiota and are derived from diet; including microbial, plant and animal sources; the microbiota itself; or from the host; with the latter mainly associated with mucins [7–9]. Mucins are heavily O-glycosylated proteins that represent the major organic components of gut epithelium glycocalyx and mucus, comprising of a central hydroxy-amino-acid rich polypeptide backbone (often referred to as the PTS repeat region for proline-threonine-serine) with O-linked glycan side chains and in gel-forming mucins N- and C-terminal cysteine rich domains that cap each end and facilitate cross-linking of the chains to form a complex mucin network [2, 10] (Fig. 1A). Central to all mucin O-glycosylated sugars is the N-acetylgalactosamine sugar (GalNAc) which links the glycan chain through an O-glycosidic bond to serine in the PTS backbone. The resulting O-glycan side chains are characterized by considerable structural diversity of GalNAc-containing core structures, backbone repeats and terminal epitopes, resulting in a highly complex and heterogeneous macromolecule [10–12] (**Fig. 1A**). Notably, only a subset of bacterial species of the gut have developed the capacity to graze on mucins, a trait thought to represent a high level of adaptation to this niche, facilitating both initial colonisation and long-term survival during absence of diet-derived glycans and thereby considered to play a key role in community development and stability [7, 9, 11, 13-18]

In order to maintain digestive and immunological homeostasis, both mucin production and grazing must be coordinated, with the microbiota contributing to both of these processes [10, 11]. Significantly, it is increasingly recognised that the outcome of unbalanced mucin grazing by the microbiota plays a key role in disease both locally, as in inflammatory bowel disease (IBD) and colonic cancer, and more distally such as in diabetes and arthritis [1, 4, 9-11, 19-21]. As a result, there has been considerable interest in recent years to improve our understanding of the mechanisms of mucin O-glycoprotein metabolism by members of the gut microbiota. The prominent human colonic microbe *B. theta* is characterized by extensive glycan degradation capabilities, a broad generalist capable of targeting plant-, microbial- and animal-derived substrates, including mucins [8, 17, 22]. Indeed, the ability of *B. theta* to access mucins has been shown to be a key trait required for effective colonisation and maintenance of this microbe in a mouse model [17]. In Bacteroidetes, the major gram-negative phylum of the gut, the glycan degrading apparatus is encoded by discrete polysaccharide utilization loci (PULs); co-regulated gene clusters that each encode a suite of cell-envelope located carbohydrate-active enzymes (CAZymes) [23] and glycan-binding and transport proteins that orchestrate the acquisition, import and degradation of a specific complex glycan [8, 24-27]. In *B. theta* 16 to 18 PULs encompassing about 100 genes are up-regulated on exposure to mucin O-glycans, either *in vitro* or *in vivo*; currently representing the largest number of PULs dedicated to a specific class of glycan[17, 28, 29]. The large number of PULs involved not only emphasises the importance of mucins utilization to *B. theta* survival, but also the complexity and heterogeneity of these substrates. In addition, only a very limited number of the induced genes have been functionally characterized[21, 30, 31] highlighting our limited understanding of mucin metabolism at mucosal surfaces.

We originally identified M60-like (M60L) peptidases as a novel subfamily of metallopeptidases significantly enriched among mucosal microbes, including *B. theta* and demonstrated that BT4244-M60L possesses metal dependent mucolytic activity in vitro [32]. Since then, several studies have investigated the structural and functional basis of BT4244-M60L O-glycopetidase activity as well as other O-glycopeptidases from M60-like and other protease families and demonstrated their application in various contexts including the detection and visualisation of mucin domain proteins, [33–37]. BT4244-M60L is part of a PUL termed PUL BT4240-50 that is up-regulated during growth of *B. theta* on mucin both *in vitro* and *in vivo* and one of five *B. theta* PULs that is important in facilitating colonisation of the mouse gut [17, 29, 38, 39]. The BT4240-50 PUL contains several other genes whose function are unknown. In a bid to enhance our understanding of the functional context of BT4244 and ultimately mucin metabolism in *B. theta*, we functionally characterized PUL BT4240-50 using a combination of genetic, biochemical, structural and cell biology approaches. We show that PUL BT4240-50 encodes a machinery that couples the processing of mucins with the metabolism of GalNAc, a core and abundant mucin O-glycan sugar. Within this machinery, various components are seen exhibit complementary binding and enzymatic activities consistent with the processing of core-1 (Galb1-3GalNAc) containing glycoproteins (Fig. 1). Genetic studies further reveal an essential role for the PUL in the competitive metabolism of mucins *in-vitro*. The data also shows that PUL BT4240-50 encodes a critical GalNAc kinase enzyme which extends the PULs metabolic function beyond mucins by virtue of its ability to phosphorylate GalNAc from diverse sources. Overall, these data support a model where GalNAc acquisition and metabolism mediate competitive growth of gut *Bacteroides* species in the presence of mucin and other GalNAc-containing macromolecules, revealing further avenues to dissect the complex mucosa-microbiota interactions in the gut and their impact on health and disease.

## Results

### Proteins encoded by the mucin activated PUL BT4240-50

The PUL BT4240-50 (Fig. 2A) is upregulated on exposure of *B. theta* to mucins both *in vitro* and *in vivo*, suggesting the encoded proteins are involved the acquisition and breakdown of a discrete structure in these complex glycoproteins [17, 38, 39]. Notably, the PUL has also been shown to be specifically upregulated during growth on core 1 disaccharide (Galβ1-3GalNAc), a core component of various mucins [17, 40] (**Fig. 1A, B**). The PUL is divided into two major operons BT4240-43 and BT4244-50 (Fig. 2A). The former operon is constitutively transcribed to a high basal level when cells are grown on glucose and up-regulated ∼11-fold (average across ORFs) in the presence of porcine gastric mucin III (PGMIII) whereas operon BT4244-50 has a lower basal level of expression and is up-regulated ∼130-fold (average across ORFs) in the same growth conditions [17]. Based on *in-silico* analyses using Pfam and KEGG annotation tools [41, 42], the PUL encodes a predicted or hypothetical protein (BT4240) with a phosphotransferase enzyme family domain, two glycoside hydrolases (GH) (BT4241 and BT4243) belonging to family GH2 and GH109 respectively, a putative inner membrane sugar transporter (BT4242), a M60-like metallopeptidase (BT4244) predicted to be surface lipoprotein and previously shown to possess O-glycopeptidase activity (Fig. 2A) [32–37] and homologues of the outer membrane (OM) canonical *Sus* proteins SusD (BT4246) and SusC (BT4247) known to be involved in cell surface glycan import [25, 26, 43, 44](Fig.2A,B). Downstream of BT4246 is the gene BT4245 predicted to be a putative surface glycan binding protein (SGBP) by virtue of the presence of a family 32 carbohydrate-binding module (CBM32) at the C-terminus as the major domain in the protein and a lipoprotein II signal (Fig. 2A). The combination of a protease and glycosidases within the same PUL suggests that this system may target both the protein and glycan moieties of mucins.

**Figure 2:**
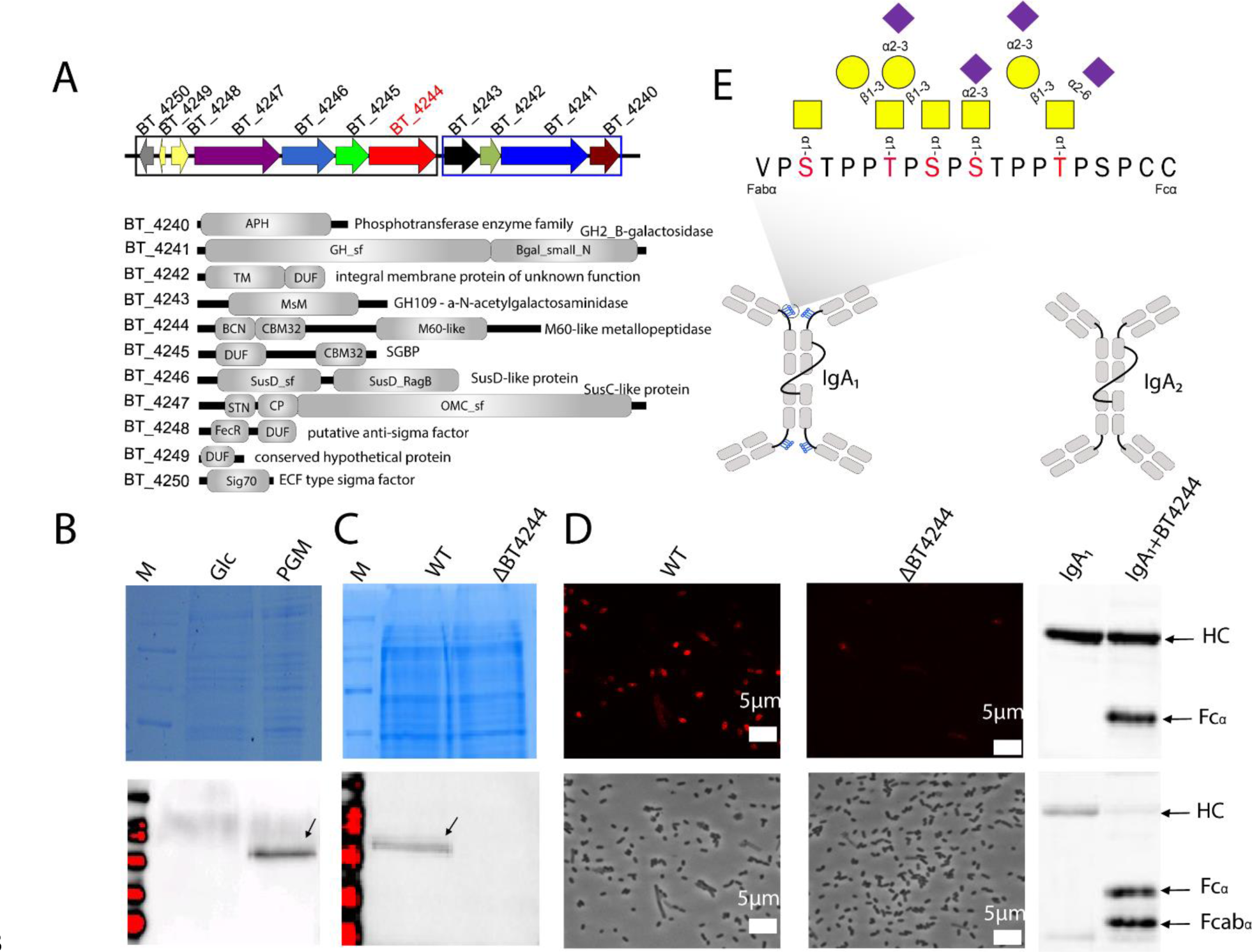
The BT4240-50 locus and cellular location and activity of BT4244-M60L glycopeptidasew. **A:** BT4240-50 locus showing genomic context of BT4244-M60L and other PUL components with predicted functions (Cazy, Pfam and KEGG annotations) [41, 42] and modular architectures. The PUL is organised into two discrete operons (boxed in black and blue). The first operon consisting BT4243-40 is expressed constitutively while the second operon contains genes encoding the hallmark SusCD surface proteins (BT4246-7), BT4245 SGBP and BT4244-M60-like, which are significantly upregulated during growth on mucins as sole carbon source [17, 32]. BT4244-M60L, is a mucin degrading metallopeptidase characterised previously [32] **B:** Induction and detection of BT4244-M60L following growth of *B. theta* in minimal medium containing either glucose (Glc) or porcine gastric mucin (PGM) as sole carbon sources. Upper panel shows Coomassie blue staining of total cell lysate while bottom panel shows western blot detection of BT4244-M60-like using anti-BT4244 antibodies raised against the purified recombinant version of the protein. **C:** Detection of BT4244 in Wild-type (WT) or BT4244 deletion strains (ΔBT4244). Upper panel shows total cell lysate stained with Coomassie, lower panel shows western blot probed with anti-BT4244 antibodies. In panels B and C the arrows indicate a band of the predicted MW of BT4244. **D:** Cellular localisation of BT4244-M60-like by antibody immunofluorescence. WT and ΔBT4244 cells were cultured in in minimal medium containing mucins and probed with same antibodies in B/D followed by fluorescently labelled secondary antibodies (upper panels - see methods). Lower panels show phase contrast image of the same cells. **E:** Proteolytic cleavage of IgA_1_ by BT4244-M60-like at the O-glycosylated hinge region. Top and bottom panels show detection of intact and cleaved IgA_1_ using anti-IgA antibodies and GalNAc-binding lectins respectively. **F:** Model structure of secretory IgA_1_ and IgA_2_ showing a O-glycosylated hinge peptide region in IgA_1_.

### The mucin-active peptidase BT4244-M60L is surface-located and cleaves the O-glycosylated peptide linker of immunoglobulin A1 (IgA_1_)

A classical functional paradigm based on the mechanisms of several previously characterised Sus-like systems depicts a key endo-acting enzyme usually located on the cell surface that cleaves the target polymer into small oligomers for import by the cognate SusC/SusD-like pair [8, 14, 25-27, 45]. The presence of a predicted type II lipoprotein signal peptide and a CBM32 domain in BT4244-M60L are indicative of a possible cell surface association [32] (Fig.2A). To investigate the expression and cellular location of BT4244-M60L, polyclonal antibodies raised against a recombinant BT4244-M60L were used to probe whole extracts of *B. theta* cells grown on glucose or PGM following western blotting. The results showed detection of a protein band with a size of ∼95kD (Fig. 2B, C), matching the expected size of BT4244-M60L. This band was absent in both glucose grown *B. theta* cells and deletion mutants (ΔBT4244) lacking BT4244-M60L (Fig. 2C). The cell surface location of the BT4244-M60L was further confirmed through immunofluorescence microscopy on fixed WT and ΔBT4244 cells grown on PGM(Fig. 2D).

We and others previously showed that BT4244-M60L is active *in-vitro* on animal mucins as well as a range of O-glycosylated proteins and model glycopeptides with a preference for Tn or Core 1 structures as the main cut sites [32–36]. However secretory IgA, like intestinal mucins, is also prominent in in the gut and the activity of BT4244 against this glycoprotein has not been tested. IgA is the most abundant immunoglobulin in mucosal secretions and shares several important features with mucins (Fig. 1A, Fig.2F) [46, 47]. Secretory IgA consists of two types including IgA_1_ and IgA_2_ (Fig. 2F)[48]. Notably, IgA_1_ contains a hinge region between its Fabα and Fcα regions which is absent in IgA_2_ (Fig.2F). The hinge region contains proline, threonine and serine repeat sequences (PTS linker) which are also O-glycosylated as in intestinal mucins, [49] thus making them potential targets for PUL BT4240-50 *in-vivo*. Incubation of BT4244-M60L with the two secretory IgA isoforms and other glycosylated/non glycosylated general protease substrates (Casein, Gelatin, Bovine serum albumin and CD44 [37, 50, 51]) followed by SDS-PAGE/Western blotting analyses with anti-IgA antibodies and GalNAc-binding lectin showed prominent degradation of IgA_1_ but not IgA_2_ (which lacks the O-glycosylated PTS linker of IgA_1_) or any of the other proteins tested (with the exception of possible slight degradation of O-glycosylated CD44; (Fig. 2E, F and Supplemental fig. 1). Overall these data reveal that IgA_1_ is a substrate for BT4244-M60L peptidase and that cleavage occurs at the O-glycosylated hinge region of the immunoglobulin.

### CBM32 domains of BT4244-M60L and BT4245-SGBP recognize galacto-configured sugars

BT4244-M60L and BT4245-SGBP are lipoproteins each containing a CBM32 and hence are likely involved in carbohydrate binding interactions at the cell surface (Fig. 2A, B). BT4244 also contains an N-terminal BACON domain with a yet to be defined function, although a previous study has suggested these are spacer domains involved in positioning the C-terminal catalytic domains on the cell surface, likely as part of their cognate SusCD utilisome[25, 52]. To gain more insights into the mucin structures targeted by BT4244-M60L and BT4245-SGBP, the ligand specificities of various modules were analysed by isothermal titration calorimetry (ITC). The N-terminal region of BT4244 containing the BACON and CBM32 domains (BACON-CBM32) was initially expressed and screened against a series of mucin-derived sugars (Table S1). BACON-CBM32 bound only to GalNAc and Gal with a preference for the amino sugar. When expressed individually and tested, it was observed that only the CBM32 domain but not BACON was capable of binding to the two sugars (Table S2; Supplemental fig 2A) consistent with a linker/spacer role earlier proposed for the BACON domain in the context of the SusCD utilisome [25, 45]. Attempts to express the CBM32 of BT4245-SGBP alone failed, however the full-length BT4245-SGBP which was successfully expressed displayed similar ligand specificity to the BT4244-CBM32, but with ∼4-fold higher affinity (Table S2; Supplemental fig 2A). Structural alignment of the CBM32 domains of BT4244, BT4245 and the *C. perfringens* GH89 enzyme (CpGH89CBM32-5, PDB: 4AAX) showed spatial conservation of key residues implicated in substrate (GalNAc) recognition in CpGH89CBM32-5 including H1392 and N1428 and F1483 corresponding to residues N352, H316 and F417 in BT4245-SGBP and H168, N202 and F260 in BT4244-CBM32 (Fig 3A, B, C). H1392 and N1428 make polar hydrogen bond interactions with the GalNAc ligand [53] in agreement with data from site directed mutagenesis and subsequent ITC analyses which showed that the equivalent residues in BT4244-CBM32 (H168, N202) are essential for binding to GalNAc (Supplemental fig. 2B). The shared specificity of the BT4244-M60L and BT4245-SGBP proteins for galactose-configured ligands is consistent with the two proteins binding O-linked glycans on the surface of *B. theta* via their CBM32 domains. Notably, an Alphafold model of the full length BT4244 protein shows that the binding site of the CBM32 domain is facing the active site of the M60-like domain, supporting a role for the CBM in either substrate targeting or positioning, or both (Supplemental fig. 2C).

**Figure 3:**
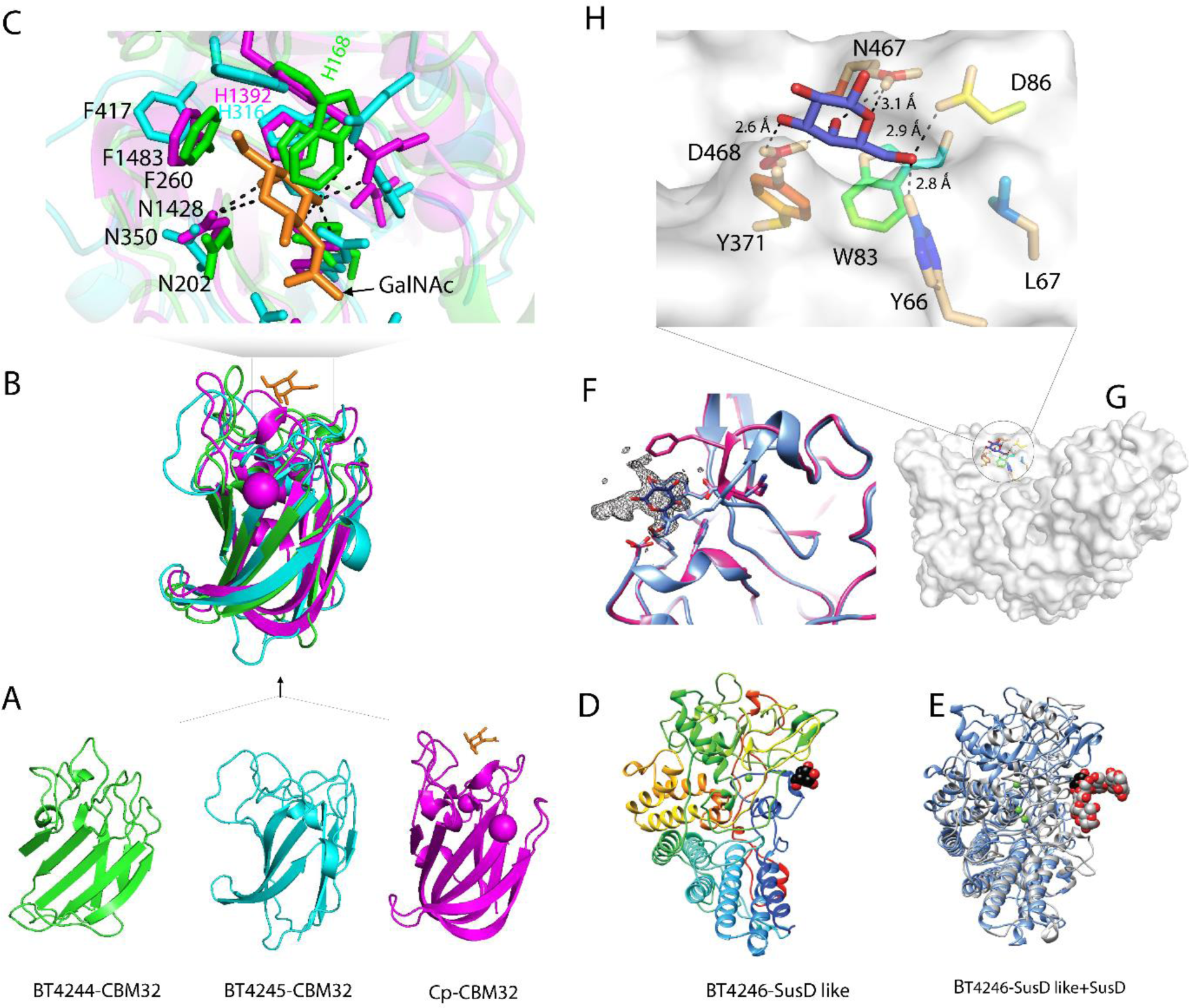
Binding site comparison of CBM32 domains from various PUL BT4240-50 encoded proteins and the crystal structure of BT4246 SusD-like protein. **A:** The BT4240-50 PUL encodes two proteins possessing CBM32 domains (BT4244-M60L and BT4245 SGBP). Alpha fold structures of both domains are compared with the crystal structure of the GalNAc binding CBM32 from a *C. perfringens* GH89 bound to GalNAc (Cp-CBM32-5, PDB: 4AAX). **B:** Alignment of all three CBM32 structures. **C:** Binding site comparison from alignment in B showing residues involved in recognition of GalNAc in CpCBM32 and their equivalents in BT4244-CBM32 and BT4245-CBM32 and hydrogen bond interactions. **D:** Crystal structure of BT4246-SusD in complex with galactose from a mucin O-glycan oligosaccharide. The protein backbone is colour-ramped blue to red from the N- to C-terminus, and a single Ca2+ is shown as a green sphere. Galactose is displayed as black and red spheres. **E:** Overlay of the canonical starch binding SusD (grey, BT3701; PDB ID 3CK9) and BT4246 (blue) with maltohexaose bound to SusD shown in grey/red and galactose bound to BT4246 shown in black/red spheres. **F:** Omit map displaying the fo-fc density (δ=3.0) for galactose in black mesh. The coordinates of the ligand-bound (blue) and semet apo (pink) structure are overlaid to demonstrate the plasticity of the loop defined by residues 462-467. **G:** Surface topology of BT4246 showing binding pocket of galactose **H:** Zoomed-in view of galactose binding site pocket of BT4246 with sugar coordinating residues displayed as sticks. Dashed lines indicate potential hydrogen-bonding interactions within 3.5Å.

### The PUL-encoded SusD-like BT4246 binds mucin-derived β-linked disaccharides

SusD-like proteins are typically surface located lipoproteins that bind specific glycan fragments generated on the surface of the cell by their PUL-encoded endo-acting enzyme and deliver these to their partner SusC-like for active import into the periplasm [14, 17, 22, 25-27, 45]. Understanding the glycan specificity of the PUL encoded BT4246-SusD-like could therefore provide further insight into the mucin components targeted by PUL BT4240-50. The ITC data showed that BT4246-SusD-like displayed quantifiable binding to the disaccharide sugars galacto-N-biose (core 1; Galβ1-3GalNAc) and lacto-N-biose (LNB; Galβ1-3GlcNAc) (with a >3-fold preference for the core 1 structure) but not the monosaccharides of the same sugars or α-linked disaccharides (Galα1-3Gal or GalNAcα1-3GalNAc) (Table S2; Supplemental fig 2A).

### Structure of BT4246-SusD-like bound to a mucin-derived oligosaccharide

The structure of BT4246-SusD-like (PDB: 5CJZ), solved to 1.8 Å, revealed the protein adopts a typical SusD-like helical fold (Fig. 3D)[54, 55]. To investigate the ligand binding site of BT4246-SusD-like, the native protein crystals were soaked in a 4% solution of O-glycan oligosaccharides purified from porcine gastric mucin (Martens 2008). In the resulting structure, unambiguous electron density was observed for a β-linked galactose, suggesting that this terminal moiety was preferentially ‘selected’ from the heterogeneous mixture. Significantly, this galactose is wedged into a shallow surface pocket located in the same place on the protein that the starch-binding site is located on SusD (rmsd 1.3Å for 151 Cα pairs, Fig. 3D, E, Supplemental fig. 2D and Table S3) [8] Additional density was observed extending off the O1 and O2 of the galactose in BT4246 but could not be definitively assigned and did not directly interact with the protein(Fig. 3F). The galactose is recognized via hydrophobic stacking with Y371 and the side chains of residues Y66, D86, R467, and D468 are proximal for hydrogen-bonding (Fig. 3G, H). Glucose cannot be accommodated in this site as the O4 would sterically clash with W83, and the hydrogen bond between the O4 and the carboxylate side chain of D468 would be lost. However, the selectivity of the protein for β-galactose is unclear, as there is no obvious steric hindrance for binding of α-linked galactose.

While there is no significant conformational change that occurred upon ligand-binding, we observed differences in orientation of the loop defined by residues 462-467 between the native protein crystals (space group P622) and that of the selenomethionine-substituted protein crystals (space group C2221) (Fig. 3F). In the selenomethionine-BT4246-SusD-like structure, F465 partially occludes the binding site, while R467, which is proximal to the galactose O1 and O5 atoms in the native crystals, is oriented out of the binding site (Fig. 3F). These changes suggest that there is some inherent plasticity of the protein surrounding the binding pocket that may facilitate ligand recognition.

### PUL BT4240-4250 encodes a β1-3-galactosidase and α-N-acetyl-galactosaminidase that target mucins and mucin-derived structures

To explore the activities of the predicted glycoside hydrolases BT4241-GH2 and BT4243-GH109, recombinant forms of the proteins were assayed against a range of PNP-monosaccharides, mucins and mucin-derived disaccharides. BT4241 was shown to be β-galactosidase cleaving several β-linked galactose containing substrates with a ∼100-fold preference for 1,3 over 1,4-linked galactose, an unusual specificity for GH2 β-galactosidases which commonly prefer the latter linkage (Fig. 4A and Table S4) [23]. The highest activity observed was against galacto-N-biose (core 1; Galβ1-3GalNAc), a structure that is known to be resistant to hydrolysis by many intestinal bacteria [11, 56] followed by lacto-N-biose (Galβ1-3GlcNAc), revealing ∼3-fold preference for GalNAc over GlcNAc at the +1 sub- site (Table S4). BT4241-GH2 was also able to release galactose residues from mucins such as PGMII and III, but only after pre-treatment with an α1,2-fucosidase to remove the fucose residues that typically cap the galactoses in these glycoproteins (Fig. 4B) [57]. The enzyme displayed no activity however against the β1,3GlcNAc in core 3 disaccharide (GlcNAcβ1-3GalNAc), a prominent core structure found in colonic mucin in particular MUC2 [58] (Fig. 4F).

**Figure 4:**
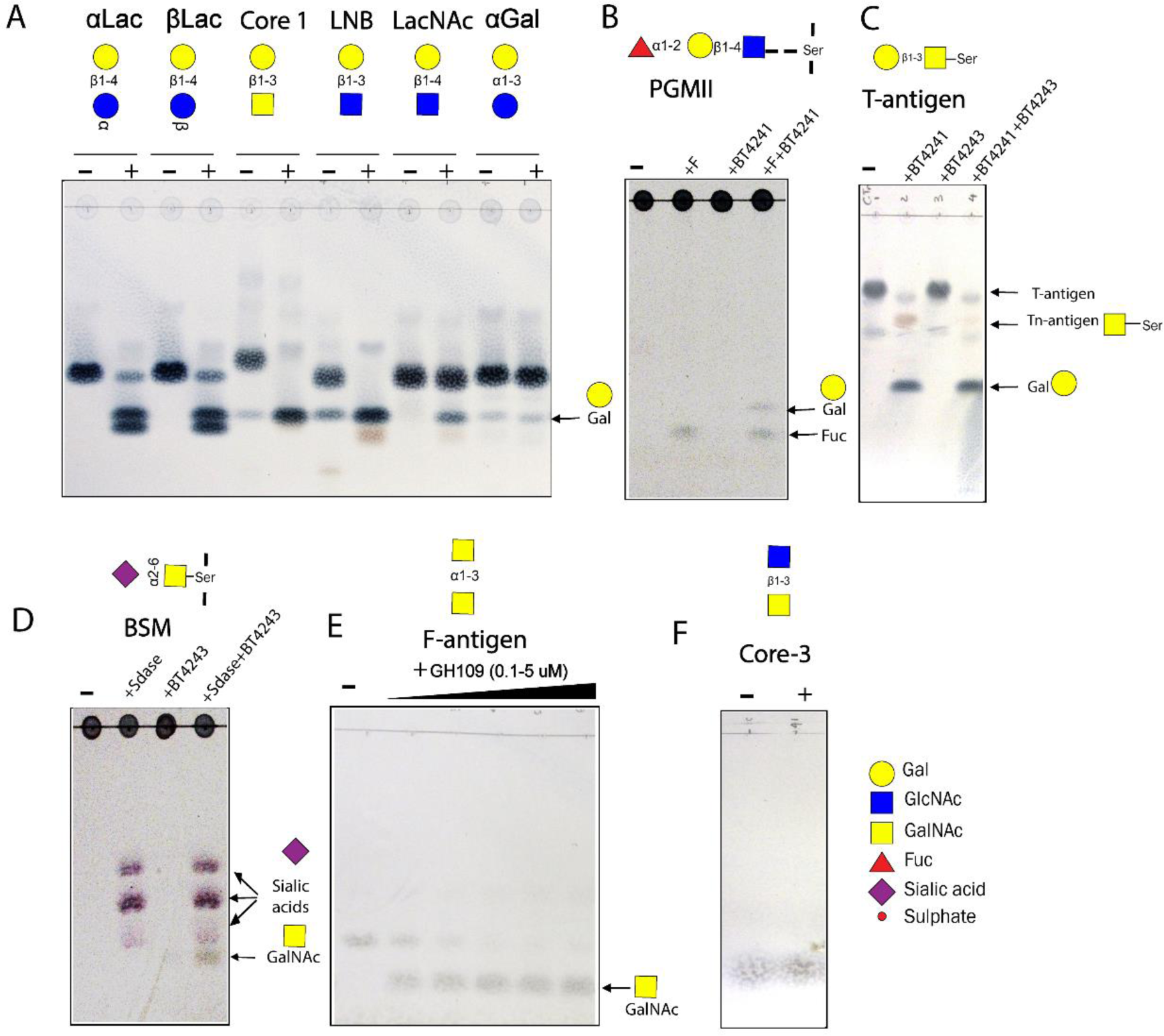
Activity of PUL BT4240-4250 encoded glycoside hydrolases against mucins and mucin-derived disaccharides. **A:** BT4241-GH2 activity against various mucin-derived disaccharides. Lane (-) represents substrate alone without BT4241-GH2 enzyme and (+) with added BT4241-GH2 enzyme. **B:** Activity of BT4241-GH2 against PGMII mucin with and without pretreatment with an α1,2-fucosidase (F). The majority of PGMII glycans contain Fucα1-2-Galβ1-4 GlcNAc structures as shown above **C:** Activity of BT4241-GH2 and BT4243-GH109 against T-antigen (Galβ1-3GalNAcα1-Ser). BT4241-GH2 releases Gal and Tn antigen (GalNAcα1-Ser) from T-antigen. BT4243-GH109 is then able to release GalNAc from Tn antigen (GalNAcα1-Ser). **D:** Activity of BT4243-GH109 against BSM. The majority of BSM glycans contain sialylated Tn antigen. After desialylation with a sialidase enzyme (Sdase) of BSM, BT4243-GH109 cleaves the uncapped GalNAc from BSM. **E:** BT4243-GH109 cleaves F-antigen (core 5; GalNAcα-1-3GalNAc) disaccharide. **F:** Treatment of core-3 with BT4241-GH2 (+ indicates with enzyme). No activity was detected against this substrate indicating the enzyme is galactose-specific.

Analysis of the specificity of the BT4243-GH109 enzyme revealed it to be an exo-acting α-N-acetylgalactosaminidase (α-GalNAc’ase) capable of hydrolysing the α1-O-linked glycosidic bond between GalNAc and serine in the Tn antigen as well as PNP-α-GalNAc (Fig. 4C and Table S4), an activity not previously described for this family. Notably, BT4243-GH109 was inactive on T-antigen (Galβ1-3GalNAcα-1Ser) unless pre-treated with the PUL-encoded BT4241-GH2 β-galactosidase (Fig 4C). The BT4243-GH109 enzyme was also active on bovine submaxillary mucin (BSM) with the amount of GalNAc released increasing significantly following pre-treatment with a sialidase enzyme. This is in agreement with a previous report showing that much of the α-GalNAc in BSM is capped with sialic acids [59] (Fig. 4D). Additionally, BT4243 was highly active on the Forsmann disaccharide sugar (GalNacα1-3GalNac) and the peripheral blood group A antigen (BgA) both of which contain a terminal α-GalNAc1-3 linkage. The Forsmann disaccharide has been reported in colonic MUC2 [58]. GH109 members have also been previously characterised and shown to have activity against BgA antigen and synthetic PNP-α-GalNAc [60–62]. Although data on their activities against Tn antigen are lacking, activity against PNP-GalNAc suggests that specificity of the GH109 is largely dictated by GalNAc and hence are also likely to be active against Tn antigen.

Other GH families with α-GalNAcase activity include GH27, GH31, GH36, GH101, and GH129. Several genes encoding these GH families except GH101 and GH129 are annotated in the genome of *B. theta* (CAZy.org). Recently, a GH31 α-GalNAcase from *B.caccae* (BACACC_01242) that targets the core GalNAc in O-glycans from fetuin and a glycopeptide, was reported [62] and a homologue of the *B. caccae* enzyme was identified in *B. theta* (BT3169) sharing significant sequence and domain features (75% ID) (Supplemental fig. 3A-C). BT3169 has also previously been implicated in O-glycan processing[55]. Analysis of BT3169-GH31 activity against Tn-antigen, BgA and desialyated BSM showed that the enzyme also exhibits α-GalNAc’ase activity, but is only active against the mucin and not Tn or BgA (Fig. 5). The lack of activity of BT3169-GH31 vs BgA is similar to the *B. caccae* enzyme and a homologue from *Enterococcus faecalis{Miyazaki, 2022 #94}*, but different to BT4243-GH109, which was active against all three substrates (Fig. 5). Notably however, BT3169-GH31 appeared to display higher activity than BT4243-GH109 against BSM, suggesting that the GH31 may play a more significant role in deglycosylating intact mucins than BT4243, which seems to prefer smaller mucin breakdown products, such as Tn antigen, as well as the terminal GalNAc decorations found in BgA. Overall, these data show that *B. theta* encodes at least two different enzymes capable of removing core GalNAc from mucins.

**Figure 5:**
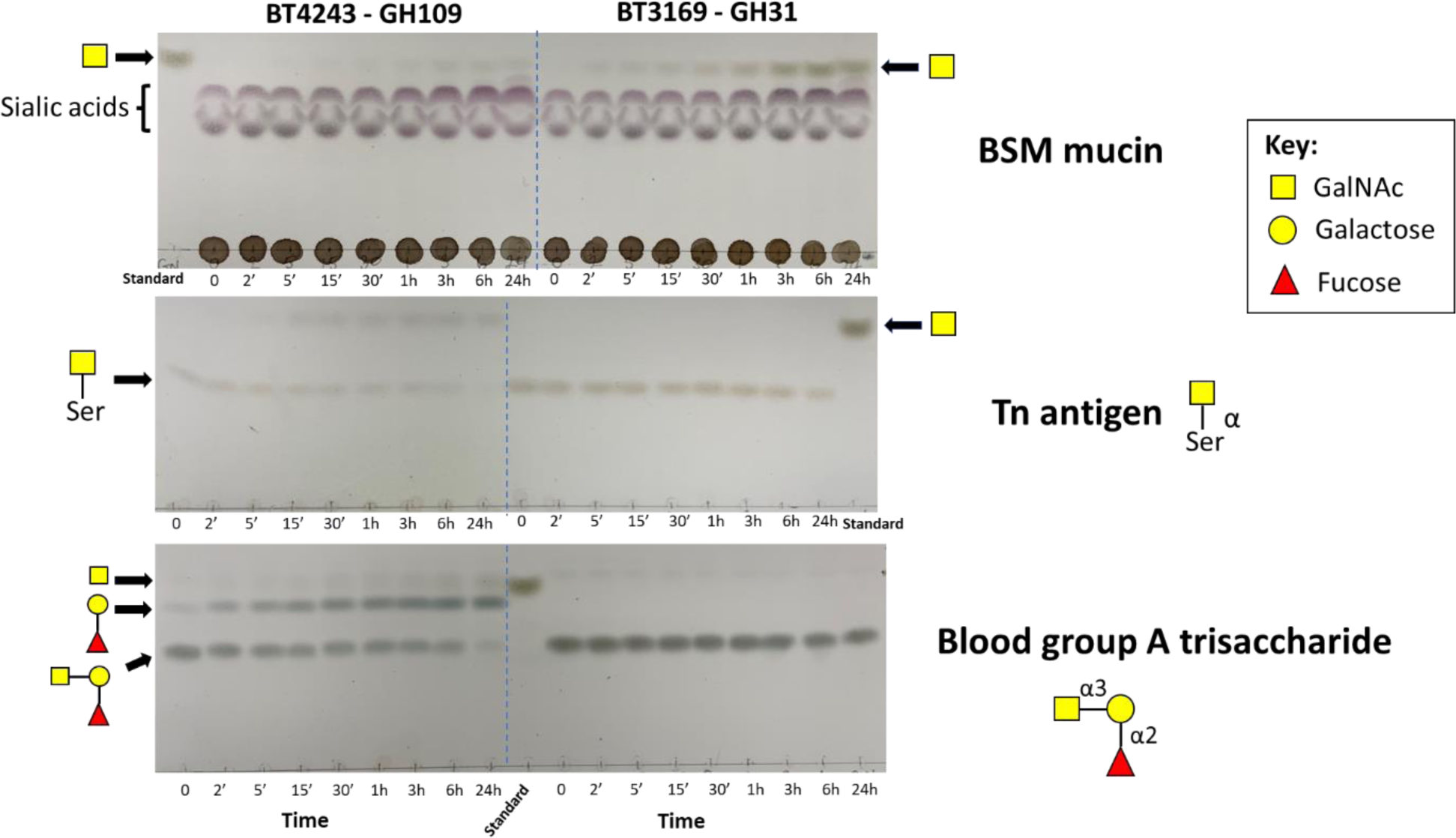
Comparison of BT4243-GH109 and BT3169-GH31 activities against mucin derived α- GalNAc containing structures. Potential substrates were incubated with the same concentration of the two α-GalNAc’ases and samples collected at various time points and analysed by TLC. BT4243-GH109 was active against all three substrates but with a strong preference for Blood group A, while BT3169-GH31 was only active against desialylated BSM although appeared to show higher activity vs the mucin than BT4243. NB. BSM was pre-treated with sialidase to uncap any potential core GalNAc residues.

### BT4240 is a GalNAc specific kinase

BT4240 is annotated as a phosphotransferase containing an APH (aminoglycoside phosphotransferase family (PF01636)) motif; Fig.2A,B) and displays 38% identity to an N-acetylhexosamine kinase from *Bifidobacterium longum* [63]. To investigate the activity of BT4240, a recombinant form of the protein was tested for its ability to phosphorylate a range of monosaccharides and hydroxy-amino acids found in O-glycans. The data revealed that BT4240 was indeed an N-acetylhexosamine kinase with a strong preference (∼275-fold) for GalNAc over GlcNAc, unlike the *B. longum* enzyme that displays similar activity against both sugars (Fig. 6, Table S4). BT4240-kinase showed no activity on any of the other monosaccharides or amino acids tested (Fig. 6A). To determine if phosphorylation of GalNAc occurred before or after release of the sugar from the peptide, the activity of BT4240-kinase against the Tn antigen (GalNAcα1-Ser) was measured in the presence and absence of BT4243-GH109 α-GalNAc’ase (Fig. 6B). The data showed that the rate of phosphorylation of GalNAc increased significantly (judging by the significant decrease in A_340nm_) following the addition of the BT4243-GH109 to the assay, indicating that phosphorylation of the sugar preferentially occurs after its release from the Tn-antigen. It may also imply that BT4240-kinase preferentially phosphorylates the sugar at position O1 which is blocked by Ser in the Tn antigen. However, the presence of the Ser may also impact phosphorylation at other sites including at position O6 which is also a popular phosphorylation site of sugars. Notably, there are no additional homologs of BT4240-kinase in the *B. theta* genome, suggesting that it may be the only GalNAc kinase encoded by the organism.

**Figure 6:**
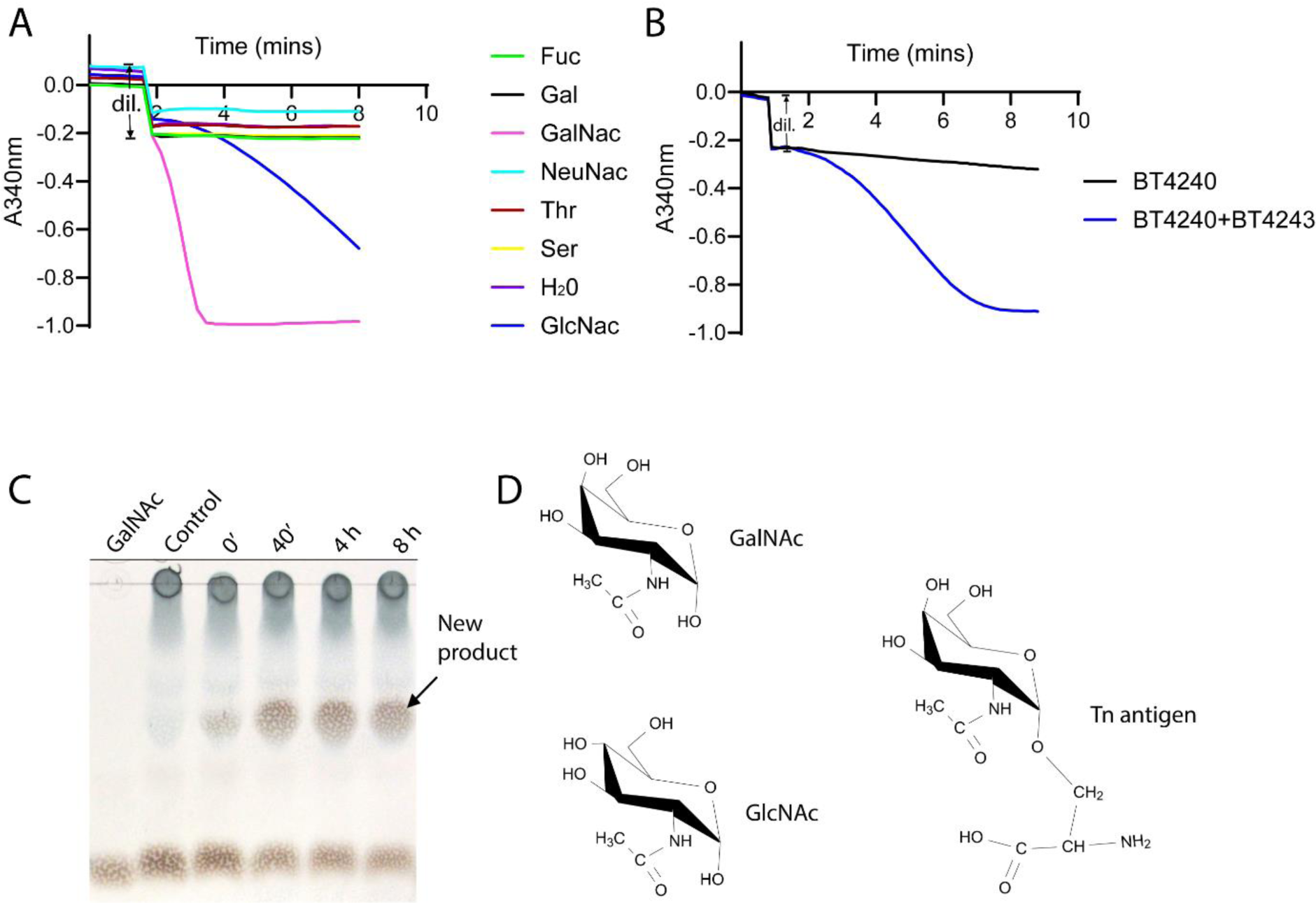
Kinase activity of BT4240 against various substrates. **A:** Kinase activity was tested against various mucin derived monosaccharides and amino acids using a linked kinase assay that generates NADH following phosphorylation of the substrate leading to a decrease in optical absorbance at 340nm (A_340_) [85]. Region of curve annotated ‘dil.’ shows an initial decrease in A_340_ due to dilution of the reaction mixture following addition of BT4240. A major decrease in A_340_ was detected for GalNAc and GlcNAc substrates indicating kinase activity. **B:** Kinase activity of BT4240-kinase on Tn-antigen and BT4243-GalNAc’ase treated Tn-antigen. **C:** TLC analysis of BT4240 kinase reaction with GalNAc showing formation of new product (likely GalNac-1-P) and depletion of GalNAc substrate over time**. D:** Structures of substrates of BT4240. NB. Intact Tn antigen shows very low activity with BT4240.

### Concerted action of PUL BT4240-50 encoded GHs provides insight into BT4244 M60-like activity against IgA_1_

The activities obtained so far for the various PUL BT4240-50 binding proteins enzymes are highly complementary and highlight a sequential substrate degradation mode culminating in the release of the core mucin O-glycan sugars GalNAc and Gal from glycoprotein substrates. As the targets for these enzymes - including the Galβ1-3GalNAc-peptide are present on a natural substrate such as IgA_1_ (Fig 7A), this offered a useful opportunity to not only assess the impact of O-glycosylation on the activity of the BT4244 M60-like glycopeptidase but also gain an insight into the hierarchical order in which the different PUL components work. To achieve this, the PUL encoded GHs and a commercially available broad-acting neuraminidase from *Clostridium perfringens* were used to sequentially de-glycosylate the antibody and at each stage treating the deglycosylated antibody with the peptidase. The results, judging by the intensity of the released Fcα fragment, showed an increase in the degradation of IgA_1_ with increasing deglycosylation up to GalNAc Fig. 7B,C). Notably, complete deglycosylation led to a significant reduction in the amount of released Fcα, evident from a 48 h time course experiment which showed a slow rate of release of the Fcα degradation fragment in a reaction containing BT4243 as opposed to one lacking the enzyme (Fig. 7B,C). These data are consistent with previous studies of BT4244 against synthetic substrates where it was demonstrated that the enzyme preferentially targets unsialylated O-glycopeptides (containing GalNAc or Galβ1-3GalNAc) and had very limited activity against the unglycosylated peptide[33]. Overall, these data support the hypothesis that BT4244-M60L is able to cleave *in vivo* glycosylated PTS repeats of IgA_1_, with both the peptide sequence and presence of GalNAc contributing to the processing rate and specificity of O-glycosylated proteins.

**Figure 7:**
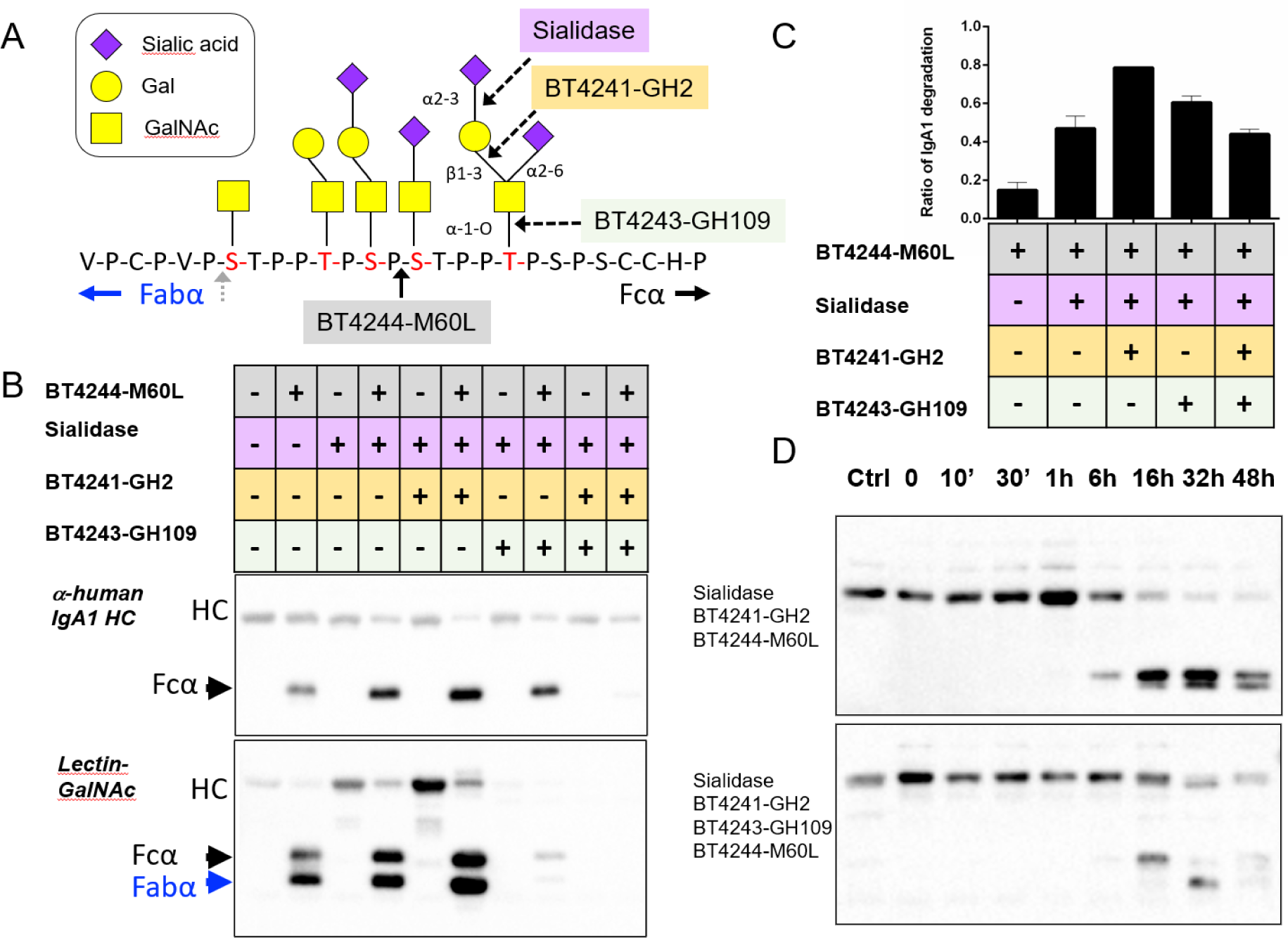
Impact of glycosylation of IgA_1_ on BT4244 M60L peptidase activity. **A:** IgA_1_ peptide hinge region showing O-glycans and target sites for PUL encoded glycoside hydrolases (BT4241-GH2 β-1,3- galactosidase and BT4243-GH109 α-GlaNAc’ase), sialidase and BT4244 M60-like peptidase. **B:** Detection of IgA_1_ breakdown products after digestion with various enzyme combinations. The upper panel shows the presence (+) or absence (-) of different enzymes in the reaction mixture with IgA_1_. Reactions were run on an SDS-PAGE gel, blotted and detected with either anti-IgA_1_ heavy chain (HC) antibodies (middle panel) or a GalNAc specific lectin (lower panel). **C:** Quantitative analyses of IgA_1_ degradation under various conditions in B **D:** Time course comparison to track the impact of the activity of BT4241 and BT4243 on BT4244-M60L activity.

### PUL BT4240-50 is important for competitive growth on mucins

To explore the importance of the PUL BT4240-50 in mucin metabolism, a deletion (or KO) mutant lacking the entire PUL BT4240-50 (ΔBT4240-50) was created and assessed for its ability to utilize porcine gastric mucin (PGM). ΔBT4240-50 and the wild type (Wt) strains grew at similar rates on glucose and in the early stages of growth on PGM (Fig. 8). The mutant however displayed a small but reproducible reduction in maximal density at the later stages of growth on PGM, suggesting a component of the PGM was not available to the mutant strain (Fig. 8). The growth defect was significantly more dramatic in competition experiments in vitro between the Wt and KO strains, where the KO was rapidly outcompeted when the carbon source was switched from glucose to PGM (Fig. 8). The data thus demonstrate that PUL BT4240-50 is important for competitive access to at least one component of mammalian mucins. To assess what this might be, we tested growth of the mutant on two of the main constituent monosaccharides of the mucin, Gal and GalNAc. Growth experiments with the monosaccharides produced an interesting result in that the PUL BT4240-50 deletion mutant completely lost its ability to grow on GalNAc but not Gal (Fig. 8), indicating that one or more components of PUL BT4240-50 is important for growth on GalNAc. Interestingly, a BT4240-kinase deletion mutant ΔBT4240 completely failed to grow on GalNAc over a 20 h period compared to the wild type strain (Fig. 8). The ΔBT4240 mutant also showed defective growth on PGMIII, similar to the levels observed for ΔBT4240-50 on the same substrate. Complementation of the ΔBT4240-50 mutant with the BT4240 also significantly restored its growth to near wild type levels. These data thus reveal that the major contribution of ΔBT4240-50 during growth on PGMIII is the metabolism of GalNAc by the kinase. This is in line with evidence showing redundancy in the activity of the other PUL BT4240-50 components such as the α-N-acetylgalactosaminidase activities observed for BT3169-GH31 and BT4243-GH109 which both release GalNAc from mucins (Fig. 5) [62] and the widespread redundancy of GH2 genes across mucin-inducible loci [19]. Additionally, a BT4244 mutant did not show a significant growth defect on PGMIII and neither was it outcompeted in competition experiments (Supplemental fig. 4) also consistent with the presence of several other predicted M60-like or proteolytic enzyme genes in the genome of *B. theta* (Table S5). In the same light, deletion of the BT4242 gene encoding the putative transporter did not affect growth on various substrates including Gal, GalNAc and PGMIII compared to the wild type strain (Supplemental fig. 4) further indicating the likelihood of redundancy in PUL BT4240-50 functions.

**Figure 8.**
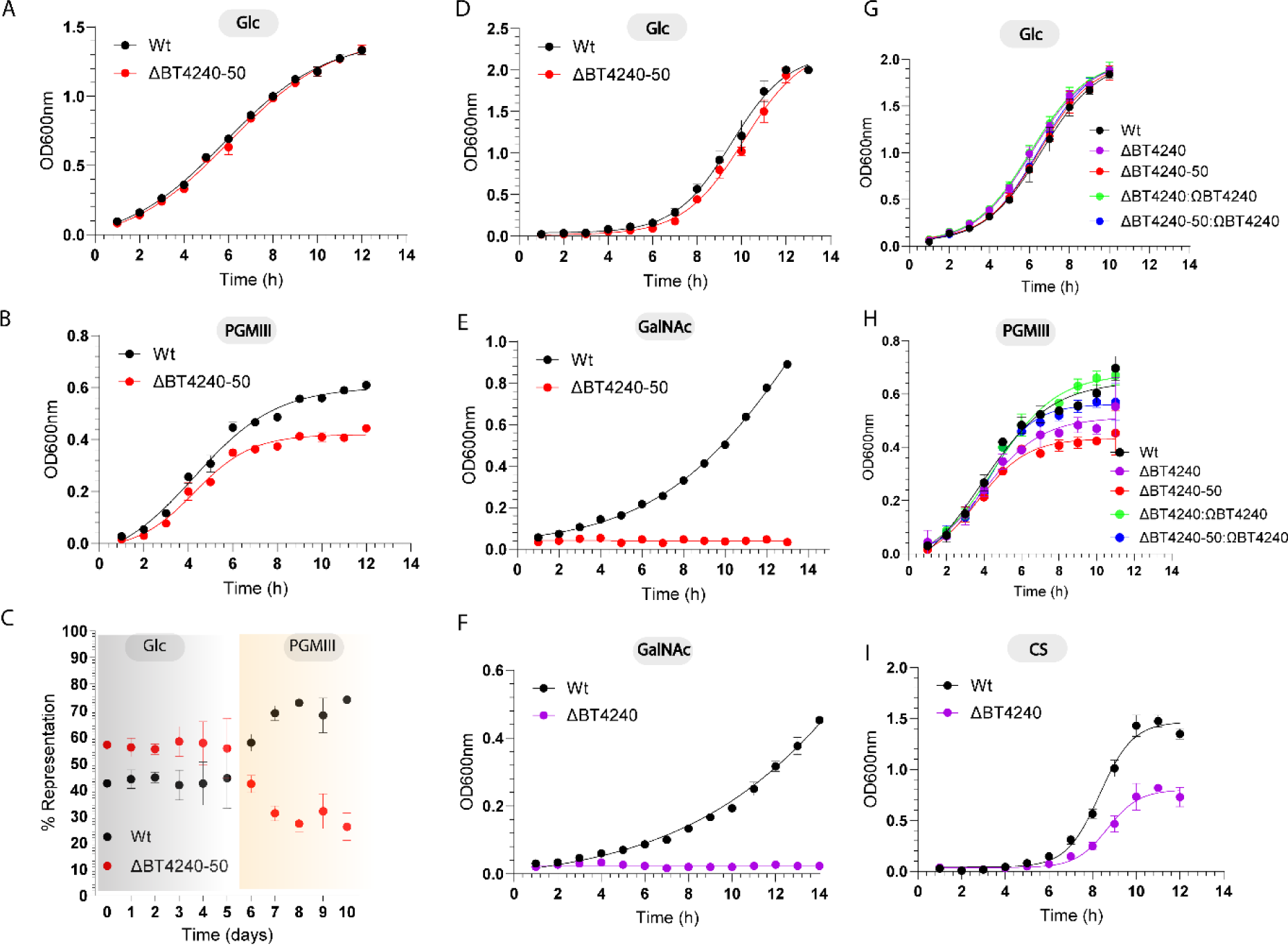
Growth and *in-vitro* competition experiments with *B. theta* wild-type and mutants on mucins, N-acetylgalactosamine and chondroitin sulphate. **A, B, D-I** shows growth profiles produced by monitoring OD600nm of each culture of deletion and complementation mutants in various nutrient sources. **C:** Shows competition experiments whereby tagged *B. theta* Wt and ΔBT4240-50 mutant cells were co-cultured in minimal medium (MM) containing Glc and then switched to MM containing PGMIII as the sole carbon source. The percentage representation of each strain was quantified by qPCR analyses. Error bars indicate standard deviations of three independent cultures. Deletion and complementation mutants are represented by Δ and Ω, respectively.

### BT4240-kinase plays a central role in global GalNAc metabolism

Out of the myriad of genes (at least 100 genes) and PULs induced during mucin metabolism and the high rate of redundancy in mucin-degrading gene families in *B. theta* [16], [27], [28], it was intriguing to observed that deletion of the BT4240-kinase alone was able to prevent growth of the organism on GalNAc. This led us to hypothesize that BT4240’s role likely extends beyond mucin metabolism. To test this, we cultured the ΔBT4240 strain in the presence of a non-mucin but GalNAc-containing glycan substrate, chondroitin sulphate (CS), which also happened to be a prominent dietary nutrient source for the human gut microbiota targeted by *B. theta* [17, 64, 65]. The mechanism of the CS PUL has been described previously, revealing that GalNAc and 5-keto, 4-deoxyuronate are major end products of the degradative process [64]. We thus anticipated that if BT4240 metabolises GalNAc from CS, the BT4240 mutant would show a significant growth defect commensurate with the proportion of GalNAc in the CS which is about 50%. This was confirmed whereby the ΔBT4240 strain was only able to grow to about 50% of the level of the wild type on CS at various stages of growth (Fig 8). The results confirm that BT4240-kinase processes GalNAc from other glycan sources like CS and likely represents a central metabolic hub for GalNAc processing in *B. theta*.

### Cellular localisation of PUL BT4240-50 proteins provides insights into the pathway for mucin breakdown in *B. theta*

To get a better understanding of the mechanism of PUL BT4240-50, the cellular locations of five PUL components including BT4240, BT4241, BT4243, BT4244 and BT4245 were investigated. To do this, the native proteins in *B. theta* were FLAG^®^-tagged at their C-termini and their locations tracked in various cellular extracts (prepared by centrifugal fractionation of whole cell lysates/sonicate of *B. theta* cells) by western blotting with antibodies against the FLAG^®^ peptide. The BT4244 protein which was earlier confirmed to be surface localised (Fig. 2B-D) was also FLAG^®^-tagged and used as control. Analyses of the lysis product of all five engineered cells showed detection of all the FLAG^®^-tagged proteins (Supplemental fig. 5). The BT4240-kinase was solely detected in the soluble fraction but not the cell membrane fractions (CMF). BT4241-GH2 was detected in both cell membrane and soluble fractions but were more enriched in the soluble fraction. BT4243-GH109 had a very similar profile to BT4241-GH2. As expected and in contrast to BT4240-kinase, BT4244-M60L was mainly enriched in the cell membrane fraction with a reduced amount in the soluble fraction. The same profile was also observed for BT4245 suggesting that the protein is also cell surface-localised as predicted for a SBGP. BT4240’s profile is in line with its *in-silico* cytoplasmic prediction. It is also interesting that BT4240 was almost completely absent from the insoluble CMF fraction as opposed to the proteins predicted to be periplasmic, which could still be detected in small amounts in the CMF fractions. It is possible that the small amounts detected in the CMF are proteins caught in transit through the inner membrane to the periplasmic space and could justify the complete absence of BT4240 in the CMF since it doesn’t have to go through the inner membrane. Combining the cellular location and activity data for each protein, a model of the function of the PUL was deduced (**Fig. 9**). In this model, naturally occurring short mucin chains (e.g., Tn and core-1-containing chains) or those generated by activity of surface glycosidases such as GH16 endoglycosidases (previously detected in *B. theta* [14]) are bound on the surface of the cell by the surface proteins BT4245-SGBP and BT4244-M60L through their respective CBM32 domains. The glycoprotein backbone is cleaved and the resulting glycopeptides imported into the cell through the PUL encoded SusCD transporter. Periplasmic enzymes BT4241-GH2 and BT4243-GH109 sequentially cleave Galβ1-3 and GalNAc1-O linkages respectively releasing Gal and GalNAc. The peptide-attached GalNAc can also be released by other GalNAcases including the BT3169-GH31. Released GalNAc from various sources (core, terminal decorations and chondroitin sulphates [64]) are imported into the cytoplasm for phosphorylation by BT4240 kinase and used for the generation of energy and carbon fixation, promoting growth (Fig. 9).

**Figure 9.**
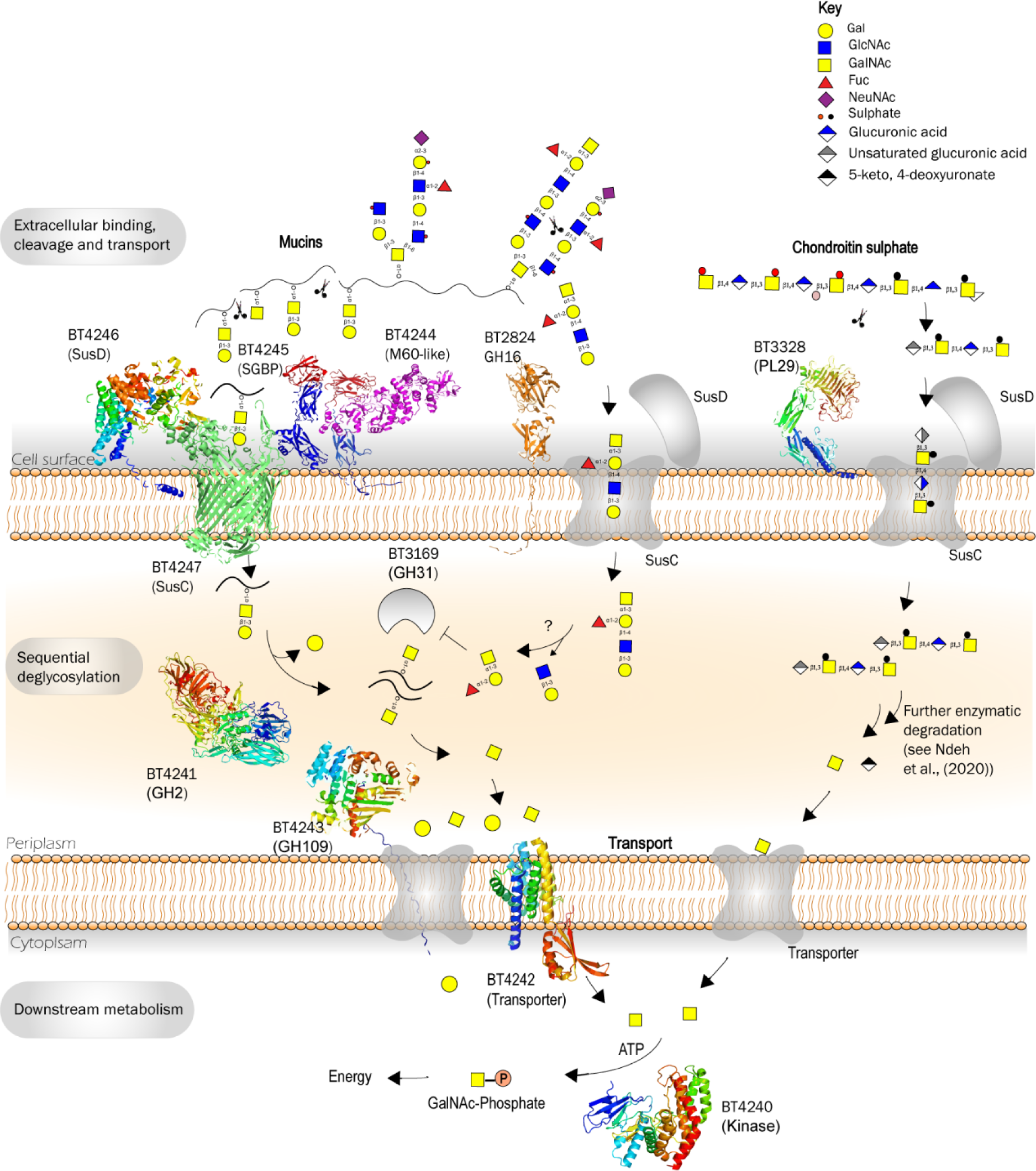
A model pathway for the breakdown of mucin O-glycans by PUL BT4240-50. Galactose- configured sugars in mucin O-glycans are bound at the cell surface by CBM32s from BT4245-SGBP and BT4244 M60L peptidase. The BT4244 M60L peptidase endolytically cleaves the glycosylated mucin peptide backbone at Tn and core 1 structures, releasing glycopeptides that are bound by the SusCD transporter proteins BT4246/BT4247 and imported across the outer membrane. GH16 endo-O-glycanases (e.g. BT2824) also cleave the mucins endolytically at the cell surface, but act on the oligosaccharide side chains, thus likely cleaving prior to the BT4144 endo-peptidase [14]. Once in the periplasm the glycopeptides (core 1 and others) are broken down to its constituent monosaccharides by the sequential action of BT4241-GH2 β1-3-galactosidase and BT4243-GH109 and BT3169-GH31 α-N-acetyl-galactosaminidases. The Gal and GalNAc released are then transported into the cytoplasm by yet to be defined or redundant transporters including BT4242, where the GalNAc is phosphorylated by the BT4240 kinase for downstream metabolism. GalNAc released from e.g. BgA decorations, as well as from chondroitin sulphate breakdown [64, 65] is also processed by BT4240 kinase.

### Origin and distribution of PUL BT4240-50 across *Bacteroides* spp. and metagenomes

To investigate the distribution of the BT4240-50 PUL across *Bacteroides* spp., a combination of BlastP searches with all proteins encoded by the PUL BT4240-50 used as query and gene neighbourhood analyses identified identical PULs to BT4240-50 in all available 13 *B. theta* annotated genomes at the IMG database [66], which include isolates from domesticated animals (Atherly and Ziemer 2014) in addition to humans (Supplemental fig. 6). Consistent with the conservation of the PUL BT4240-50 between human and animals *B. theta* strains, all 41 *B. theta* reconstructed genomes from human faecal metagenomes [67] contained the PUL. These analyses also identified BT4240-50-like PULs in the annotated genomes from *Bacteroides faecis* and *B. caccae* (Supplemental fig. 7). Scanning 287 human faecal metagenome sequence data identified the PUL BT4240-50 in 54% of the samples, a similar level to the *B. theta* α-mannan PUL2 BT3773-92(56%) (Supplemental fig. 8) [24]. The combined distribution of the three similar PULs; *B. theta* BT4240-50, *B. faecis* KCYDRAFT_01771-81 and *B. caccae* BACCAC_01835-46, was found across 73% of the human gut metagenomes analyzed, suggesting that the encoded proteins play an important role in microbial survival in the gut (Supplemental fig. 8) [24, 25].

The operon BT4240-43 is also found among different *Bacteroides* species including those without homologs of the operon BT4244-47 in the direct neighbourhood, or without operons with BT4244- M60L homologs altogether (operon BT4244-47) (Supplemental fig. 8). This suggests that both operons evolved independently from each other and are only known to be combined into one PUL in *B. theta*, *B. faecis* and *B. caccae*. For PULs from other *Bacteroides* spp. possessing BT4244-47-like gene configurations, alternative predicted CAZyme associations where identified, suggesting targeting of different glycoproteins/glycan structures to those described for PUL BT4240-50 (Supplemental fig. 8).

## Discussion

The ability to utilize mucins as a nutrient source is a key trait shared by a subset of the eubiotic (health promoting) colonic microbiota, which include representatives from all known gut phyla, that contributes to both mucosal homeostasis and community resilience [7, 9, 11, 13-18]. Despite the importance of mucolysis to human health, our understanding of the mechanisms of mucin breakdown by the microbiota is still very limited. This is contributed by several factors including the complexity and diversity of mucin glycans. Members of the Bacteroidetes which are one of the most abundant microorganisms in the human gut microbiota encode glycan-degrading systems called PULs which enable them to capture and metabolise a diverse range of glycans. Typically one or more of these loci are dedicated per glycan, however for mucins, over 16 to 18 PULs encompassing about 100 genes are up-regulated on exposure to mucin O-glycans, representing the largest number of PULs and genes dedicated to a specific class of glycan [17, 28, 29]. To date, only a very limited number of the PUL genes have been functionally characterized [14, 30, 31, 68-72] and it is not known to date if all 16-18 mucin PULs are essential for mucin metabolism. Previously we and others have characterised the founding member BT4244-M60L of a novel family of proteases termed M60-like capable of degrading mucins [32–36]. BT4244-M60L’s association with PUL BT4240-50 suggests that the enzyme acts in concert with other enzymes, therefore it was expected that the characterisation of the PUL BT4240-50 would not only shed new light into the mechanism of a mucin inducible PUL but also contextualise the proteolytic activity BT4244-M60L. Our data from the biochemical characterisation of various PUL BT4240-50 components suggest that it is capable of metabolising mucin glycans containing the core-1 sugar Galβ1-3GalNAc, as well as the core GalNAc (Tn antigen). Significantly, our combined cell localisation, functional and comparative genomic data support a model where extracellular cleavage of the mucin peptide backbone is coordinated with uptake of core-1-containing glycopeptides and their subsequent deglycosylation in the periplasm. The pathway shares similarities with the classical PUL paradigm whereby a large glycan is endolytically cleaved by extracellular enzymes to facilitate capture and import into the periplasmic space for further processing, however, while glycanases are generally widely implicated in this activity, the use of proteases in this case is consistent with the nature of the target substrate, in this case a glycoprotein reflecting the versatility and adaptability of *B. theta* to different nutrient conditions. It is also worth noting that surface exposed endo-acting O-glycanases in family GH16 have been reported [14] implying that more than one surface endo-acting enzyme is deployed by *B. theta* to target different components of intestinal mucins (Fig. 9).

Secreted proteases targeting non-glycosylated terminal domains of mucins are generally recognized as key virulence factors for a variety of mucosal pathogens, allowing disruption of the cross-linked mucin chain network to facilitate access to the epithelium [5, 10, 49]. Restricted targeting of the terminal domains is thought to be due to the heavy O-glycosylation of the central PTS rich region protects it from proteolysis. By contrast, BT4244-M60L and other M60-like glycopeptidases encoded by the microbiota are able to cleave the glycosylated PTS backbone of the intestinal mucins and may provide an explanation for the report that very little intact mucin is found in faeces [5], a fact that is at odds with an exclusively extracellular glycosidase model of mucin breakdown, which would result in the peptide backbone remaining largely intact. The capacity to fully process mucins substrates by members of the microbiota also ensures the recycling of an important fraction of the large biomass of secreted mucins a process which contributes to gut-microbiota homeostasis both in terms of energy and as a carbon source for the microbiota and enterocytes through short chain fatty acid production [73]. We also demonstrate that BT4244-M60L can cleave the O-glycosylated peptide linker of IgA_1_, providing new insights into the complex mechanisms of glycoprotein processing by the microbiota. Notably, BT4244 preferentially targets PTS with Tn-antigen and core 1 decorations in IgA1 and does not efficiently cleave sialyted structures, consistent with the recent data from several other studies looking at 4244 specificity [33]. IgA_1_ is also a prominent mucosal surface glycoprotein and has been shown to be targeted by several protease enzymes. Degradation of IgA_1_ is thought to represent an immune evasion mechanism in these microbes. Interestingly, BT4244 cleaves at the same site as a *Prevotella melaninogenica* protease and the pathogenic microbes *Haemophilus aegyptius*, HI-1: *Haemophilus influenzae* [74, 75]. Whether this PUL enables *B. theta* to extract glycans from IgA_1_ or evade immune action is not known and the functional significance of its IgA_1_ protease activity is yet to be established. The successful exploitation of IgA_1_ in understanding BT4244 activity however, demonstrates that it could serve as a useful alternative or proxy substrate in place of synthetic glycopeptides for screening and analysing glycoprotease enzymes [33, 76], especially those from the M60-like protease family where most members remain uncharacterised to date.

Genetic studies of PUL BT4240-50 demonstrated a critical role played by the PUL in host and animal derived glycoprotein metabolism particularly in providing competitive access to the prominent mucin and chondroitin sulphate sugar GalNAc. This is supported by *in-vitro* competition or fitness data using porcine gastric mucins as carbon source, inability of the deletion mutant ΔPUL BT4240-50 and Δ BT4240 to grow on GalNAc as sole carbon source and various complementation experiments showing that BT4240 kinase enzyme is capable of significantly restoring the growth defects caused by the deletion of PUL BT4240-50. This also implies that the majority of the other PUL components exhibit redundant activities during growth on porcine mucins *in-vitro,* which is not surprising given the significant expansion of mucin PULs and GH families in the *B. theta* genome and data showing that BT3169-GH31 was also capable of extracting core GalNAc from mucins. It was however very interesting that despite the expansion of mucin PULs, a single enzyme BT4240-kinase was responsible for the metabolism of such a critical and core component of mucins as GalNAc. Indeed, all mucin-O-glycans contain a GalNAc anchor, implying that BT4240 is likely a key player for competitive colonisation *in-vivo*. An even more intriguing observation was the involvement of BT4240 in CS-derived GalNAc metabolism reflecting a probable role in global GalNAc metabolism. Indeed a mutant lacking BT4240 was unable to grow in minimal medium with GalNAc as the sole carbon source while the same mutant showed about a 50% reduction in growth on CS reflecting not only the proportion of the GalNAc but also suggesting that GalNAc metabolism is completely halted in the mutant.

While the characterization of PUL BT4240-50 provides new insight into the mechanism of mucin breakdown by the microbiota, it is notably only one piece of a rather large and complex puzzle with several PULs containing multiple putative CAZymes, sulfatases and proteases that are activated during growth on mucin O-glycans [8, 17, 28]. These findings highlight both the complexity and heterogeneity of the substrate (mucins and other sources of glycans) and the fact that unlike most glycans, the breakdown of mucins appears to require the cooperative action of multiple PULs, each likely targeting a discrete structure. Indeed, for PUL BT4240-50 to access its target the core substrate in complex highly branched mucin regions, it would first require significant ‘debranching’ of the heterogeneous glycan side chains of mucin (Fig. 1) by other mucin-sensitive PULs and also, likely other members of the mucolytic microbiota. Similarly, it is possible that components of the PUL BT4240-50 are utilized by *B. theta* to access substrates other than core-1. For instance, as the GalNAc-Ser/Thr structure is conserved in all mucin cores [40] (Fig. 1), the BT4243-GH109 (and BT3169-GH31) could also be used by *B. theta* to release GalNAc from other core structures once the capping structures had been removed by other enzymes from mucin targeting PULs. Furthermore, kinase BT4240-kinase appears unique in Bt, suggesting it is essential for the metabolism of this monosaccharide derived from various sources such as other core structures, terminal epitopes or even other types of glycan such as CS (which is made of ∼50% GalNAc), and known to be metabolised by *B. theta* [22, 64, 77] and bacterial lipopolysaccharides [78, 79]. These observations could explain the higher basal level expression of operon the operon BT4240-43 (Fig. 2A) [17].

It is worth noting that core 1 structures though present, are relatively rare in human MUC2, the major colonic mucin, which mainly comprises core 3, and to a lesser extent core 2 and 4, suggesting other *B. theta* PULs play a more significant role in the degradation of colonic MUC2 [58, 80]. Interestingly, among the *B. theta* PULs induced by host O-glycans there are an additional three candidate surface proteases which could cooperate with BT4244-M60L to degrade different parts of the MUC2 polypeptide backbone and/or other mucin types (Table S5). Notably core 1 is more common in secreted mucins from the upper digestive tract (e.g. MUC5B from saliva and MUC5AC and MUC6 from the stomach and small intestine) and thus PUL BT4240-50 could play an important role in the breakdown of these non-colonic mucins as they converge to the colon, somewhat mirroring exposure of *B. theta* to dietary glycans. Similarly IgA_1_, with mucin-like PTS repeats decorated with core 1 [48] which was shown here to be a substrate for BT4244-M60L, is by far the more abundant isotype of secretory IgA in the upper part of the digestive (and respiratory) tracts (60%-95%), with the colon being the only site of the digestive tract where the non-O-glycosylated IgA_2_ is more abundant than IgA_1_ (∼60:40) [81]

Notably, 73% of scanned human gut metagenomes possess at least one of the three identified BT4240- 50-like PULs (Supplemental fig. 8), supporting the general importance of this apparatus for gut survival. Furthermore, the absolute conservation of PUL BT4240-50 across all *B. theta* isolates from human and animals gut is also consistent with the locus playing a central role in the biology of *B. theta*, contrasting with PULs with a more ephemeral status such as those targeting diet-derived plant glycans [67] or a fungal derived glycan targeting PUL [24]

To conclude we have functionally characterized one of several uncharacterised mucin inducible PULs PUL BT4240-50 from the genome of the prominent human gut microbe *B. theta,* revealing key mechanistic insights into how it coordinates the processing of mucin and other glycoprotein structures. Critically, we have demonstrated that PUL BT4240-50 is essential for *in-vitro* competitive growth on mucin glycans by *B. theta* and encodes a key kinase enzyme which makes a major contribution to mucin and GalNAc metabolism by phosphorylating the latter and enabling its utilisation as an energy or carbon source or both, promoting competitive growth. These data advances our knowledge of the vital metabolic processes that govern host-microbiota interactions at mucosal surfaces and highlights GalNAc as a key metabolite mediating microbiota-gut interactions.

## Acknowledgments

We would like to thank Dr D. Bulmer (Newcastle University) for assistance with microscopy and Dr A. Zhu (European Molecular Biology Laboratory) for providing the annotation of the 41 reconstructed *B. theta* genomes from human faecal metagenomes. We also thank Carl Morland for expert technical assistance, Claire Sawyers for technical assistance and Nicholas Pudlo and Prof Eric Martens from the University of Michigan Medical School for the generous donation of mucin O-glycan oligosaccharides for co-crystallization with BT4246.

## Materials and Methods

### Sources of carbohydrates

Porcine gastric mucin (PGM) II and III, bovine submaxillary mucin (BSM), LacNAc, Lacto-N-biose, core 1 disaccharide (galacto-N-biose), α-Galactobiose, monosaccharides and PNP-linked monosaccharides were from Sigma. T-antigen (Galβ1,3GalNAcα1-Ser) and Tn-antigen (GalNAcα1-Ser) were purchased from Dextra Laboratories (Reading, UK) and PNP- GlcNAcβ1,3GalNAc (PNP-core 3) from Santa Cruz Biotechnology (Dallas, Texas, USA).

### Cloning, expression and purification of recombinant proteins

The genes encoding the mature forms (i.e. lacking their predicted signal sequences) of the BT4240-4250 proteins characterized in this study were amplified from *B. theta* VPI-5482 genomic DNA using the primers shown in **Table S6**. The PCR products were cloned into pET28 (Novagen) or pRSETA (ThermoFisher) expression vectors and sequenced to ensure the fidelity of the cloning process. Recombinant plasmids were transformed into BL21 (DE3) or BL21 (DE3) Tuner cells (Novagen) and were incubated in 1L LB in 2L conical flasks at 37°C, 180 rpm until OD_600_ ∼0.6. The cultures were then cooled to 16°C before addition of 1 mM IPTG (final) to induce recombinant gene expression and incubation at 16°C, 180rpm overnight. Cells were harvested by centrifugation (3000*g*, 4°C), resuspended in Talon buffer (20 mM Tris-HCl, pH 8.0, 100 mM NaCl) and lysed by sonication. Recombinant His-tagged protein was purified from cell free extracts in a single step using immobilized affinity chromatography (IMAC) with Talon resin (ClonTech) and dialyzed overnight into 20 mM Tris-HCl, pH 8.0. The amount of purified protein was quantified using the predicted molar extinction coefficient as determined by the ProtParam tool (Expasy server). Proteins were used fresh or stored in aliquots at −20°C until required.

### Growth and genetic manipulation of *B. theta*

*B. theta* was routinely cultured under anaerobic conditions in 5 ml of TYG (tryptone-yeast extract-glucose medium) or minimal medium (MM) containing 0.5-1% (w/v) of an appropriate carbon source plus 1.2 mg/ml porcine hematin (Sigma) as described previously [22]. Growth of cultures was monitored using a Biochrom WPA cell density meter (Cambridge, UK).

*B. theta* knockout strains where created using counter-selectable allelic exchange as described previously [82]. Signature-tagged strains of wild type and ΔBT4240_50 *B. theta* mutant used in competition experiments were created and selected by integration of unique 24bp tag sequences into one of two *att* sites in the genome of *B. theta* following conjugation with an S17-1 *λ pir E.coli* strain harboring an NBU-2 based vector (pNBU2-tetQb) as described previously [17]. Selection of integrants was done on BHI-agar containing gentamycin (200 mg/ml) and tetracycline (1 mg/ml). Genomic DNA was extracted from single colonies after culturing in TYG using the GenElute™ Bacterial Genomic DNA kit (Sigma) and screened for tag insertions by PCR using primers flanking various *att* sites. A list of all primers used in various genetic manipulations is provided in **Table S6**.

### In vitro competition of wild type and ΔBT4240-50 *B. theta* strains on mucin

Competition experiments were carried out to evaluate the contribution of the BT4240-50 locus to *B. theta* fitness on PGMIII. Approximately equal amounts of the signature-tagged wild type and ΔBT4240-50 deletion mutant initially grown in TYG were mixed and 100 µl used to inoculate minimal medium containing 1% glucose (MM-Glc). After overnight growth, 100 µl of culture was used to inoculate fresh MM-Glc media every day for 5 days before switching to MM-PGMIII (1% w/v) for a further 5 days. Genomic DNA was extracted from 2 ml of overnight culture from each time point using the GenElute™ Bacterial Genomic DNA kit (Sigma) and 10 ng analysed by qPCR using a Light Cycler 480 real-time PCR system (Roche) to quantify the proportion of each tagged strain.

### Cellular localization studies

The expression and cellular localization of BT4244 M60L was determined using rabbit polyclonal antibodies (Eurogentec) generated against a purified recombinant version of the protein lacking the lipoprotein signal sequence. *B. theta* cells grown on MM-containing PGMII to mid exponential phase (OD_600_ ∼0.4) where fixed in an equal volume of 9% formalin in PBS, pH 7.4, by rocking gently for 90 min at 25°C. Cells were washed twice with PBS then pelleted by centrifugation for 5 min at 2400*g* and re-suspended in 1 ml of PBS. This washing process was repeated twice before incubation with blocking solution (1ml of 2% normal goat serum, 0.02% NaN_3_ in PBS) overnight at 4°C. Cells were then centrifuged again at 17000*g* for 1 min and blocking solution discarded. For labelling, cells were incubated with 0.5 ml of a 1/1000 dilution (in blocking solution) of rabbit polyclonal anti-BT4244 antibodies for 2 h at 25°C. Cells were washed again as above by centrifugation at 17000*g* for 1 min. Detection of bound antibodies was carried out using a 1/200 dilution of Alexa- Fluor® 594 conjugated goat anti-Rabbit IgG secondary antibodies (Molecular Probes) by incubating for 1 h at 25°C in the dark. Cells were then pelleted and washed with PBS as above. To 100 µl of cells, one drop (∼50 µl) of ProLong Gold anti-fade reagent (Life Technologies) was added and labelled bacterial cells were mounted onto glass slides for imaging. Phase contrast and fluorescence images were captured using an Andor iXonEM+ 885 EMCCD camera coupled to a Nikon Ti-E microscope.

FLAG^®^ tagging of the C-terminus of BT4240-4250 encoded proteins was done by incorporating the FLAG^®^ peptide (DYKDDDDK) DNA sequence into the genome of *B. theta* using the primers listed in **Table S6**. Strains containing FLAG^®^-tagged proteins were grown overnight in 10ml cultures MM- PGMIII (1% w/v), harvested by centrifugation (2400*g*, 10 min, 4°C), washed in PBS buffer and disrupted by sonication for 2 min on ice. This was followed by the separation of membrane and soluble fractions using ultracentrifugation as described previously ^21^. Fractions were analysed by Western blotting using a 1/10,000 dilution of primary rabbit anti-FLAG^®^ antibodies (Eurogentec) followed by a 1/5000 dilution of secondary donkey anti-rabbit IgG conjugated to HRP (Santa Cruz biotechnology, USA).

### Glycoside hydrolase and sugar kinase assays

#### Thin layer chromatography (TLC)

TLC analysis of sugars was performed using silica TLC plates (Sigma) as previously described [65]. Reactions (3-6 µl of each) were spotted on to plates and samples resolved in butanol/acetic acid/water buffer (2:1:1) in glass tanks. Plates were then air-dried and dipped in either orcinol/sulfuric acid (0.5% (w/v) orcinol in 10% sulfuric acid) or diphenylamine–aniline– phosphoric acid (DPA, used to detect sialic acid [83]) developers before being dried at 70°C for up to 10 min to visualize sugars. Reactions (50 µl total) were set up in 1.5 ml microcentrifuge tubes containing enzyme (BT4241 or BT4243 at 0.3-1.2 µM) and substrate (5-10 mM disaccharides or 4 mg/ml mucins) in 20 mM Tris-HCl, pH 7.5 and incubated at 37°C for 1 h. The sialidase used for pretreatment of BSM was from *Arthrobacter ureafaciens* (Calbiochem) and the fucosidase used for pretreatment of PGM was from *Bifidobacterium bifidum* [84].

#### Enzyme kinetics

The kinetic parameters of the BT4241 GH2 β-galactosidase against disaccharide substrates was determined by the continuous monitoring of galactose release using a D-Gal detection kit (Megazyme International) at 37°C in 20 mM MOPS buffer, pH 7.0. For core 1 and LNB disaccharides reactions contained 0.3 µM of BT4241-GH2 and a range of substrate concentrations from 0.05-1.1mM. For LacNAc, 3 µM of enzyme was used with substrate concentrations ranging from 0.4- 6.0mM. To screen enzymes against paranitrophenol (PNP)-linked monosaccharides, 1 µM enzyme (final) was added to 0.5 mM substrate and PNP release determined at 37°C in 20 mM MOPS buffer, pH 7.0, by continuously monitoring the increase in A_400_. For BT4243-GH109 kinetics, 0.4 or 0.7µM enzyme was added to substrate (PNP-α-GalNAc or PNP-β-GalNAc, respectively) at a range of concentrations from 0.03-0.5 mM. BT4240 kinase activity was determined using a coupled enzyme ATPase assay as described previously [85]. BT4240 (between 0.1-0.5 μM final) was added to a range of monosaccharides or amino acids (between 0.2-50 mM) at 37°C in 20 mM MOPS buffer, pH 7.0 and the reaction rate monitored by the decrease in A_340_ from the conversion of NADH to NAD. Graphs of initial rate versus substrate concentration were generated and fit to the Michaelis–Menten equation using GraphPad Prism v6.0 to determine enzyme kinetic parameters.

### IgA cleavage assay

Human myeloma IgA_1_ or IgA_2_ isoforms (Calbiochem), at 0.33 mg/ml were separately incubated with 1 µM of recombinant BT4244-M60L peptidase and incubated at 37°C in a water bath for 16h. Samples were then analysed by SDS-PAGE followed by Western blotting. For Western blotting, PVDF membranes containing blotted proteins were washed in excess PBS-Tween 20 for 1h at room temperature before application of a 1:2000 dilution of primary mouse Anti-Human IgA_1_ antibodies conjugated to biotin (SouthernBiotech). Blots were washed 10 min 3 x in PBS followed by treatment with a 1:2000 dilution of ExtrAvidin®−Peroxidase conjugate (Sigma) in washing solution for 1h. A final washing step was performed as above before chemiluminescence detection with luminol/enhancer solutions from the Biorad Immun-Star™ Western C™ Chemiluminescence kit (Bio-rad) and Luminol and enhancer solutions were mixed in equal proportions (1 ml each), spread on washed membranes and chemiluminescent signals detected using a ChemiDoc XRS system (Bio-rad). To determine the effect of deglycosylation of IgA_1_ on BT4244 peptidase activity, IgA_1_ was pre-treated with a sialidase from *Clostridium perfringens* (Sigma), BT4241-GH2 β-galactosidase (0.5 µM) and BT4243-GH109 α-N-acetylgalactosaminidase (1 µM) enzymes in different combinations overnight in a final volume of 15 µl in 20 mM Tris-HCl, pH 8.0, 100 mM NaCl, before incubation with BT4244- M60L peptidase (0.7 µM) for 16h. IgA deglycosylation was monitored using a biotin–conjugated *Helix aspersa* agglutinin (HAA) (Sigma) that binds specifically to GalNAc. This was performed on separate blots containing replicate samples and using a 1:1000 dilution of the lectin.

### N-terminal sequencing of BT4244-M60L cleaved IgA_1_

IgA_1_ (15 μg) was digested with 1 µM BT4244 for 16 hr at 37 °C in a final volume of 30 µl and run on an SDS-PAGE gel. Digested proteins were then transferred to PVDF membranes and stained for 2 h with freshly made Coomassie blue solution (0.4% (w/v) Coomassie blue, 20% methanol, 10% glacial acetic acid). The membrane containing transferred proteins was washed in 50% methanol solution containing 10% glacial acetic acid and the two cleavage products excised using a scalpel blade into 1.5 ml microcentrifuge tubes. Samples were sent to Alphalyse (Denmark) for N-terminal Edman sequencing.

### Isothermal titration calorimetry

ITC was performed using a MicroCal VP-ITC machine. Titrations (25 x 10 µl injections) were carried out in 20 mM Tris-HCl buffer, pH 7.5 at 25 °C. The reaction cell contained protein at 100 µM, while the syringe contained ligand (between 10-50 mM). Integrated heats, minus control heats of dilution, were fit to a single site binding model using Origin v7.0 to derive *K*_a_.

### Cloning and Expression of BT4246 in *E. coli* for crystallization

The gene fragment corresponding to BT4246 lacking its endogenous signal peptide (residues 25 - 642) was amplified from *B. theta* genomic DNA using the primers shown in **Table S6**. The gene product was ligated into a modified version of pET-28a (EMD Biosciences) containing a tobacco etch virus (TEV) protease recognition site. The pET28-BT4246 plasmid was transformed into Rosetta (DE3) pLysS cells (EMD Biosciences). Transformed cells were grown at 37°C for 20 h, and then the plates were scraped to inoculate culture media for protein expression. For native protein expression, the cells were grown in 1L of TB / Kan50 / Cm20 (in 2L baffled flasks) at 37°C until they reached an OD_600_ ∼0.6, when the temperature was turned down to 22°C. Approximately 30 min after lowering the temperature, IPTG was added to a final concentration of 0.5mM, and the cells continued to grow overnight (∼16h). Cells were harvested by centrifugation at 6,000*g*, and the cell pellets were stored at −80 °C until protein purification. Selenomethionine substituted protein was produced using a premade SelenoMet expression medium (Molecular Dimensions). Cells were grown for 16h in 100 ml Minimal Media starter cultures containing 40 μg/ml methionine. Cells were then pelleted at 6,000*g* and washed three time with sterile H_2_O then transferred to 1 x 2 litre baffled flasks containing the SelenoMet expression media supplemented with 40 μg/ml of selenomethionine. The cells were grown at 37°C until they reached an OD_600_ ∼0.6, then the temperature was lowered to 22°C and the cells were grown for an additional 16 h. Cultures were pelleted via centrifugation and stored at −80°C prior to purification.

### Purification of native and SeMet-substituted BT4246

Native and selenomethionine-substituted BT4246 proteins were purified using a 5 ml Hi-Trap metal affinity cartridge (GE Healthcare) according to the manufacturer’s instructions. The cell lysate was applied to the column in His Buffer (25 mM NaH_2_PO_4_, 500 mM NaCl, 20 mM imidazole, 1 mM TCEP pH 7.5) and proteins were eluted with an imidazole gradient (20–300 mM). The His-tag was removed by incubation with recombinant TEV (1:100 molar ratio of TEV to protein) at room temperature for 3 h, then for 16 h at 4°C while dialyzing against His Buffer. The cleaved protein was re-purified on the 5 ml Ni column to remove undigested target protein, the cleaved His-tag and His-tagged rTEV. Purified proteins were dialyzed against 20 mM HEPES, 100 mM NaCl pH 7.0 prior to crystallization, and concentrated using Vivaspin 15 (10,000 MWCO) centrifugal concentrators (Vivaproducts, Inc.).

### Crystallization and Data Collection

Crystallization conditions were screened via the hanging drop method of vapor diffusion in 96-well plates and using Hampton Screen kits (Hampton Research). Native crystals were obtained at 4°C via hanging drop vapor diffusion from protein concentrated to 8.8 mg/ml and mixed 1:1 with the well solution containing 22-24% PEG 3350 and 150 mM CsCl. To obtain a ligand-bound structure, we soaked these native crystals in a 4% solution of oligosaccharides derived from porcine gastric mucin, prepared by alkaline borohydride treatment as previously described[17]. Semethionine-substituted proteins crystals were obtained at 20°C as hanging drop experiments using 22.56 mg/ml protein against a well solution of 22-24% PEG 3350 and 200 mM KCl, using grains of SiO_2_ in the hanging drop to drive nucleation. These crystals were then micro-seeded onto a hanging drop containing 9.92 mg/ml of SeMet substituted protein and SiO_2_ against a well solution of 23% PEG 3350 and 200 mM KCl. Crystals that nucleated half way off of the SiO_2_ were then transferred to a hanging drop containing 17.28 mg/ml of SeMet substituted protein, against a well solution of 20% PEG 3350 and 200 mM KCl. All crystals were serially transferred into a cryoprotectant composed of 80% crystallization media, 20% ethylene glycol and flash-frozen in liquid nitrogen prior to data collection. X-ray diffraction data were collected at the Life Sciences Collaborative Access Team (LS-CAT) at the Advanced Photon Source at Argonne National Labs, Argonne, IL. X-ray data were processed with HKL2000 and scaled with SCALEPACK [86]. The structure was determined from the SAD data using the AutoSol subroutine within the Phenix software package [56, 87]. The selenomethionine-substituted model of BT4246 was then utilized for molecular replacement in Phaser [88] against the native X-ray data sets. The final models for all structures were derived from iterative cycles of manual model building in Coot [89] and refinement in Phenix. Data collection and refinement statistics are reported in **Table S2**.

### Bioinformatic analyses

Metagenomic analyses of the occurrence of the BT4240-50 PUL and selected PULs for comparison form *B. theta* and other *Bacteroides* spp. in human gut metagenome data was performed as previously described [24, 25]. Briefly PUL DNA sequences were used as query in BLAST searches (word size of 11) and hits were recorded when they passed the following cutoff criteria: E values <1E-20, and nucleotide identities >90% over a length > 100 bp. A PUL was considered present in a given sample when at least 2 distinct BLAST hits were recorded over the length of the query sequence. The details for the query sequences are as follows: *B. theta*VPI-5482, PUL *BT_4240-50*, gb|AE015928.1|, positions 5582950-5601300, 18,350 bp. *Bacteroides faecis* MAJ27, PUL *KCY_RS0108600-KCY_RS0108650* (*BT4240-50-*like), ref|NZ_AGDG01000020.1|, positions 51630- 69946, 18,316 bp. Bacteroides caccae ATCC 43185, PUL *BACCAC_01835-46* (*BT4240-50-*like), B_caccae-MSIQ_Cont1328, gb|AAVM02000003.1|, positions 306812-328036, 21,224 bp. *Bacteroides caccae* ATCC 43185, inulin PUL *BACCAC_02727-31*, gb|AAVM02000006.1|, positions 41899-59372, 17,473 bp. *Bacteroides plebeius* DSM 17135, porphyran PUL *BACPLE_01672-708*, gb|ABQC02000019.1|, positions 132998-78523, 54,476 bp. *B. theta* VPI-5482, α-mannan PUL1 *BT_2620-32*, gb|AE015928.1|, positions 3262908-3277966, 15,059 bp. *B. theta* VPI-5482, α-mannan PUL2 *BT_3773-92*,

gb|AE015928.1|, position 4893415-4928241, 34,827 bp. *Bacteroides uniformis* ATCC 8492, xyloglucan PUL1 *BACUNI_00315-26*, gb|AAYH02000032.1|, positions 43453-71885, 28,433 bp.

