## Supplemental material for "A genetic locus in the gut microbe *Bacteroides thetaiotaomicron* encodes activities consistent with mucin-O-glycoprotein processing and plays a critical role in *N*-acetylgalactosamine metabolism"

**Supplemental data**

**Table S1. ITC data for BT4244 BACON-CBM32.**

|  | Sugar | Ka × 10^3^ (M^-1^) |
| --- | --- | --- |
| Fuc (10mM) | | NB^a^ |
| Gal (50mM) | | 0.40 (±0.03) |
| GalNAc (20mM, 40mM) | | 0.16 (±0.01) |
| GlcNAc (20mM) | | NB |
| NeuAc (10mM) | | NB |
| Man (20mM) | | NB |
| Xyl (40mM) | | NB |
| Gal-6S (2.5mM) | | NB |
| Fucα1-2Gal (2mM) | | NB |
| GalNAcα1-Ser (2.5mM) | | TLTFc |
| Galβ1-3GalNAc (Core 1; 10mM) | | 0.20 (±0.01) |
| Galα1-4Glc (lactose; 20mM, 50mM) | | 0.23 (±0.00) |
| Galβ1-4GlcNAc (10mM) | | TLTF^b^ |
| Galβ1-3GlcNAc (10mM) | | TLTF^b^ |
| Fucα1-2Galβ1-4GlcNAc (2.5mM) | | NB |
| BSM (0.1%) | | NB |
| PGMIII (0.1%) | | NB |

^a^NB: no binding detected. ^b^TLTF: weak binding observed but affinity too low to fit (Ka <~0.1 × 10^3^ M^-1^).

**Table S2. ITC data for BT4244 and BT4245 CBM32 domains and BT4246 SusD-like binding to mucin derived mono- and di-saccharides.**

| Protein | Sugar | Ka × 10^3^ (M^-1^) |
| --- | --- | --- |
| BT4244-CBM32^a^ | GalNAc | 0.40 (±0.03) |
|  | Gal | 0.16 (±0.01) |
|  | NeuAc | NB^b^ |
|  | Fuc | NB |
|  | GlcNAc | NB |
|  | Lactose | 0.23 (±0.00) |
|  | Core 1 | 0.20 (±0.01) |
| BT4245 (SGBP^c^) | GalNAc | 1.20 (±0.2) |
|  | Gal | TLTF^d^ |
|  | NeuAc | NB |
|  | Fuc | NB |
|  | GlcNAc | NB |
|  | Lactose | 0.21 (±0.02) |
|  | Core 1 | 0.36 (±0.01) |
|  | F antigen | 0.88 (±0.06) |
| BT4246 (SusD-like) | GalNAc | TLTF |
|  | Gal | TLTF |
|  | NeuAc | NB |
|  | Fuc | NB |
|  | GlcNAc | NB |
|  | α-Galactobiose | TLTF |
|  | Lactose | TLTF |
|  | LacNAc | TLTF |
|  | LNB | 0.13 (±0.07) |
|  | Core 1 | 0.45 (±0.03) |
| BT4244-BACON |  |  |
|  | GalNAc | NB |
|  | Gal | NB |
|  | NeuAc | NB |
|  | Fuc | NB |
|  | GlcNAc | NB |
|  | α-Galactobiose | NB |
|  | Lactose | NB |
|  | LacNAc | NB |
|  | LNB | NB |
|  | Core 1 | NB |

^a^ CBM32 alone tested. ^b^NB: no binding detected. ^c^SGBP: surface glycan binding protein. Full length protein lacking signal peptide was tested. ^d^TLTF: weak binding observed but affinity too low to fit (Ka <~0.1 × 10^3^ M^-1^). Lactose: Galβ1-4Glc, LacNAc:: Galβ1-4GlcNAc, LNB: Galβ1-3GlcNAc Core 1: Galβ1-3GalNAc. α-Galactobiose: Galα1-3Gal, F-antigen (core 5): GalNAcα1-3GalNAc.

**Table S3. Data Collection and Refinement Statistics for BT4246 SusD-like.**

|  | SeMet | Native | Nat/O-glycan |
| --- | --- | --- | --- |
| PDB id | 5CK1 | 5CK0 | 5CJZ |
| Resolution (Å) | 31.83 - 1.835 (1.901 - 1.835) | 46.23 - 1.996 (2.068 - 1.996) | 41.88 - 1.803 (1.867 - 1.803) |
| Space group | C 2 2 21 | P 6 2 2 | P 6 2 2 |
| Unit cell (a,b,c) | 110.7, 115.7, 116.6 | 156.2, 156.2, 114.7 | 155.9, 155.9, 114.3 |
| Total reflections | 447332 (35783) | 264067 (24638) | 975782 (80697) |
| Unique reflections | 65430 (6379) | 55908 (5331) | 74235 (7283) |
| Multiplicity | 6.8 (5.6) | 4.7 (4.6) | 13.1 (11.1) |
| Completeness (%) | 99.66 (98.08) | 99.25 (96.63) | 98.25 (98.13) |
| Mean I/sigma(I) | 37.41 (13.30) | 8.03 (1.66) | 10.95 (1.03) |
| Wilson B-factor | 16.93 | 16 | 21.06 |
| R-merge | 0.1095 (0.213) | 0.2025 (0.881) | 0.2301 (1.797) |
| R-meas | 0.1185 | 0.228 | 0.2392 |
| CC1/2 | 0.986 (0.959) | 0.963 (0.315) | 0.99 (0.458) |
| CC* | 0.996 (0.99) | 0.991 (0.692) | 0.997 (0.793) |
| R-work | 0.1671 (0.1981) | 0.1867 (0.2507) | 0.1712 (0.2970) |
| R-free | 0.2013 (0.2595) | 0.2265 (0.3068) | 0.2050 (0.3568) |
| non-hydrogen atoms | 5584 | 5565 | 5811 |
| macromolecules | 4752 | 4861 | 4861 |
| ligands | 13 | 5 | 30 |
| water | 819 | 699 | 920 |
| Protein residues | 585 | 603 | 603 |
| RMS(bonds) | 0.006 | 0.007 | 0.007 |
| RMS(angles) | 0.93 | 0.97 | 0.92 |
| Ramachandran favored (%) | 97 | 97 | 97 |
| Ramachandran outliers (%) | 0.17 | 0.33 | 0.17 |
| Clashscore | 1.68 | 1.91 | 1.43 |
| Average B-factor | 19.6 | 13.2 | 19.2 |
| macromolecules | 17.8 | 12.2 | 17.3 |
| ligands | 21.4 | 25.1 | 25.5 |
| solvent | 30.4 | 19.4 | 28.9 |

**Table S4: BT4241-GH2, BT4243-GH109 and BT4240-kinase kinetic data.**

| Enzyme | Substrate | *k*_cat_  (s^-1^) | *K*_m_  (mM) | k_cat_/*K*_m_  (s^-1^ mM^-1^) |
| --- | --- | --- | --- | --- |
| BT4241-GH2^a^ | Core 1  (Galβ1-3GalNAc) | 14.2  (±0.6) | 0.36  (±0.04) | 40.0 |
|  | LNB  (Galβ1-3GlcNAc) | 6.1  (±0.6) | 0.5  (±0.1) | 12.2 |
|  | LacNAc  (Galβ1-4GlcNAc) | 0.18  (±0.02) | 1.4  (±0.3) | 0.13 |
| BT4243-GH109 | PNP-α-GalNAc | 7.8  (±0.4) | 0.03  (±0.005) | 260 |
|  | PNP-β-GalNAc^b^ | 1.9  (±0.1) | 0.08  (±0.01) | 23.7 |
| BT4240-kinase | GalNAc | 3.0  (±0.04) | 0.9  (±0.04) | 3.3 |
|  | GlcNAc | 0.34  (±0.03) | 27.5  (±4.8) | 0.012 |

^a^Both GH enzymes were initially screened against a range of PNP-linked monosaccharides (β-Gal, β-Glc, β-Man, α-GalNAc, β-GalNAc, α-GlcNAc, β-GlcNAc, β-L-Fuc, α-L-Fuc, α-Gal and α-Glc).

Activity for BT4241-GH2 was only observed with pNP-β-Gal.

Activity for BT4243-GH109 was only observed with PNP-α-GalNAc and PNP-β-GalNAc.

^b^GH109 enzymes have previously been shown to act on both α and β linked substrates due to their unusual NAD-dependent hydrolysis mechanism (55).

**Table S5. Additional candidate surface proteases from *B. theta* PULs induced by mucin.**

| Locus tag of predicted protease | Merops family | Predicted PUL^a^ | LipoP prediction^b^ |
| --- | --- | --- | --- |
| BT0212 | Subfamily S8 unassigned peptidase (MER028054) | BT0206-14 | SpII score=17.8 margin=11.9 |
| BT3015 | M60L | BT3012-15 | SpII score=20.5 margin=2.3 |
| BT3960 | Subfamily C2A unassigned peptidase (MER003948) | BT3958-61 | SpII score=13.7 margin=12.0 |

^a^ *B. theta* PULs upregulated during growth on mucins *in vitro* or *in vivo* (17).

^b^ LipoP 1.0 server [90].

**Table S6. Primers used in this study.**

| Name | Protein id/genetic modification | RE | Sequence (5’-3’) |
| --- | --- | --- | --- |
| Primers for cloning into *E. coli* expression vectors |  |  |  |
| BT4240for | Kinase | BamHI | CGCGGGATCCATGAAAGATTTATCAAGTATTGTAGC |
| BT4240rev | Kinase | XhoI | CCGGCTCGAG TTATCCATTAACCAAGCACTCATTG |
| BT4241for | GH2 | BamHI | CGCGGGATCC ATGGCCGAAAAGACATCCGACAA |
| BT4241rev | GH2 | XhoI | CCGGCTCGAG TTATCGATAATCATATTTGGCGGC |
| BT4243for | GH109 | BamHI | CGCGGGATCC CAAAAGACAAAAGCAAAGTTCTCT |
| BT4243rev | GH109 | XhoI | CCGGCTCGAG TTATTCGGCAAAAGCATGTCTGTA |
| BT4244-FLfor | M60L-full length | BamHI | CGCG GGATCC AAG GAT ACC GAA AAA TCG ATT ATA |
| BT4244-FLrev | M60L-full length | EcoRI | CCGG GAATTC TTA TAA CAG AAT ACG TTT TCC GTC |
| BT4245for | SGBP | NcoI | CGCGCCATGGACAATTATGACGATACCTATCC |
| BT4245rev | SGBP | XhoI | CCGGCTCGAGTTCGGACAGTATGAACAGACT |
| BT4246for | SusD-like | Nhe1 | GCGATCGCTAGCGATTATCTAGACGTCGTTCCACC |
| BT4246rev | SusD-like | Xho1 | GCGATCCTCGAGTTAATATCCGGGTGCTTGTACC |
| BT4244-CBM32for | CBM32 from M60L | NcoI | CGCG CCATGG ACATCAAGGTTACACCAACC |
| BT4244-CBM32rev | CBM32 from M60L | XhoI | CCGG CTCGAG CGTATTTGTTTTGTAAAATTCCATTTC |
| BT4244-BCNfor | BACON domain from M60L | BamHI | CGCG GGATCC AAG GAT ACC GAA AAA TCG ATT ATA |
| BT4244-BCNrev | BACON domain from M60L | EcoRI | CCGG GAATTC TTAGCCTCCGGTTGGTGTAAC |
| BT4244-BCN CBM32 for | BACON-CBM32 | BamHI | CGCG GGATCC AAG GAT ACC GAA AAA TCG ATT ATA |
| BT4244-BCN-CBM32rev | BACON-CBM32 | EcoRI | CCGG GAATTC TTA CGT ATT TGT TTT GTA AAA TTC CAT TTC |
| Primers for genetic manipulation of Bt |  |  |  |
| ∆4240-50F1for | BT4240-50 deletion | BamHI | CGCGGGATCCGGATCGATTTCAAGCATAACAAAT |
| ∆4240-50F1rev | BT4240-50 deletion | None | ATGGGTATGAATACGTTTGAGCCTTGGAGAATGGAAAATGGATAATTAGT |
| ∆4240-50F2for | BT4240-50 deletion | None | ACTAATTATCCATTTTCCATTCTCCAAGGCTCAAACGTATTCATACCCAT |
| ∆4240-50F2rev | BT4240-50 deletion | XbaI | CGCGTCTAGAGTATCTTCATTTACCACAGTATGGAT |
| ∆BT4244F1for | BT4244 deletion | BamHI | CGCGGGATCCCATGAGTTGCATATACATCACCAC |
| ∆BT4244F1rev | BT4244 deletion | None | TCTTAAAAACTAATACAGCAAAAAAGAATAATTTAACTTAACACACATTAC |
| ∆BT4244F2for | BT4244 deletion | None | GTAATGTGTGTTAAGTTAAATTATTCTTTTTTGCTGTATTAGTTTTTAAGA |
| ∆BT4244F2rev | BT4244 deletion | XbaI | CGCGTCTAGACTCTTGAAGAGGGACAAACGTG |
| BT4240-flagF1 | Flag-tagging BT4240 | BamHI | CGCGGGATCCATGAAAGATTTATCAAGTATTGTAGC |
| BT4240-flagF2 |  | None | gactacaaagacgatgacgacaaaTAATGGAGAATGGAAAATGGATAATTAG |
| BT4240-flagR1 |  | None | TTATTTGTCGTCATCGTCTTTGTAGTCTCCATTAACCAAGCACTCATTG |
| BT4240-flagR2 |  | XbaI | CGCGTCTAGATCAGCGGATTGTATGACAAGTTTC |
| BT4241-flagF1 | Flag-tagging BT4241 | BamHI | CGCGGGATCCATTACAAACTCAAACCCAAGGAG |
| BT4241-flagF2 |  | None | gactacaaagacgatgacgacaaaTAACCTTAGTATAAATTTTAAAGAAAAGAC |
| BT4241-flagR1 |  | None | TTAtttgtcgtcatcgtctttgtagtcTCGATAATCATATTTGGCGGCTT |
| BT4241-flagR2 |  | XbaI | CGCGTCTAGACTACGCTTTGAAGTAGTTTGAAC |
| BT4243-flagF1 | Flag-tagging BT4243 | BamHI | CGCGGGATCCACTCAGATTGTAGCTTTATGCG |
| BT4243-flagF2 |  | None | gactacaaagacgatgacgacaaaTAAGCCATTATACCTTATTAATATATAAAC |
| BT4243-flagR1 |  | None | TTAtttgtcgtcatcgtctttgtagtcTTCGGCAAAAGCATGTCTGTA |
| BT4243-flagR2 |  | XbaI | CGCGGCGGCCGCCAGTAATAGGTGATATGATCTGTAT |
| BT4245-flagF1 | Flag-tagging BT4245 | BamHI | CGCGGGATCCGACAATTATGACGATACCTATCC |
| BT4245-flagF2 |  | None | gactacaaagacgatgacgacaaaTAATAAATGAGAGACAAAGGGTGAGT |
| BT4245-flagR1 |  | None | TTAtttgtcgtcatcgtctttgtagtcTTCGGACAGTATGAACAGACTAAA |
| BT4245-flagR2 |  | XbaI | CGCGTCTAGATGTTGACTCCGGGATTTAGCATA |
| Tag1 for | Signature tag-1 | None | ATGTCGCCAATTGTCACTTTCTCA |
| Tag11 for | Signature tag-11 | None | ATGCCGCGGATTTATTGGAAGAAG |
| Tag rev |  | None | CACAATATGAGCAACAAGGAATCC |
| NBU2att1for |  | None | CCTTTGCACCGCTTTCAACG |
| NBU2att1rev |  | None | TCAACTAAACATGAGATACTAGC |

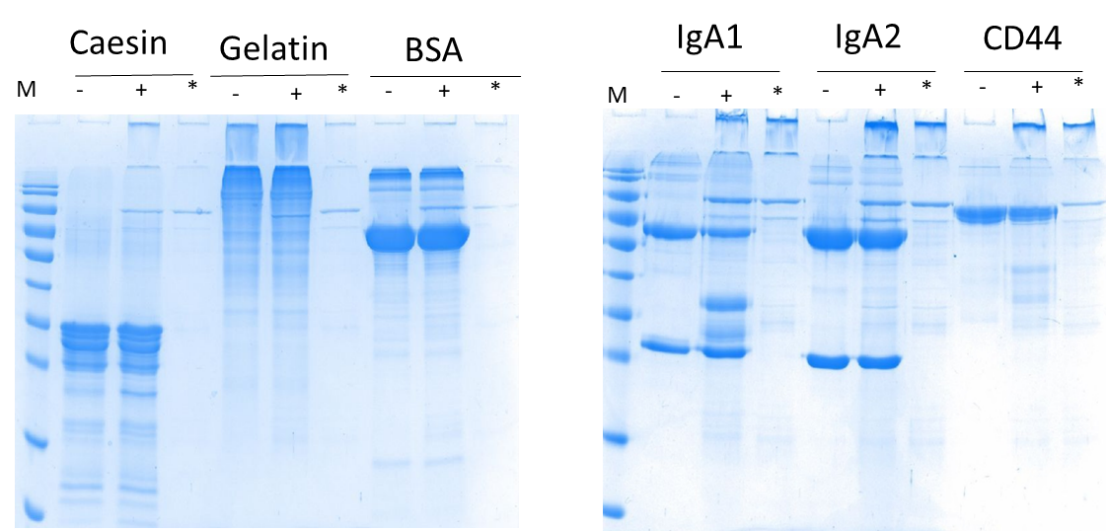

**Supplemental Figure 1**: **Activity of BT4244-M60-like against potential protein and glycoprotein substrates.** **-/**+ signs represent putative substrate without and with addition of BT4244 enzyme. * is for recombinant BT4244 enzyme alone without substrate.

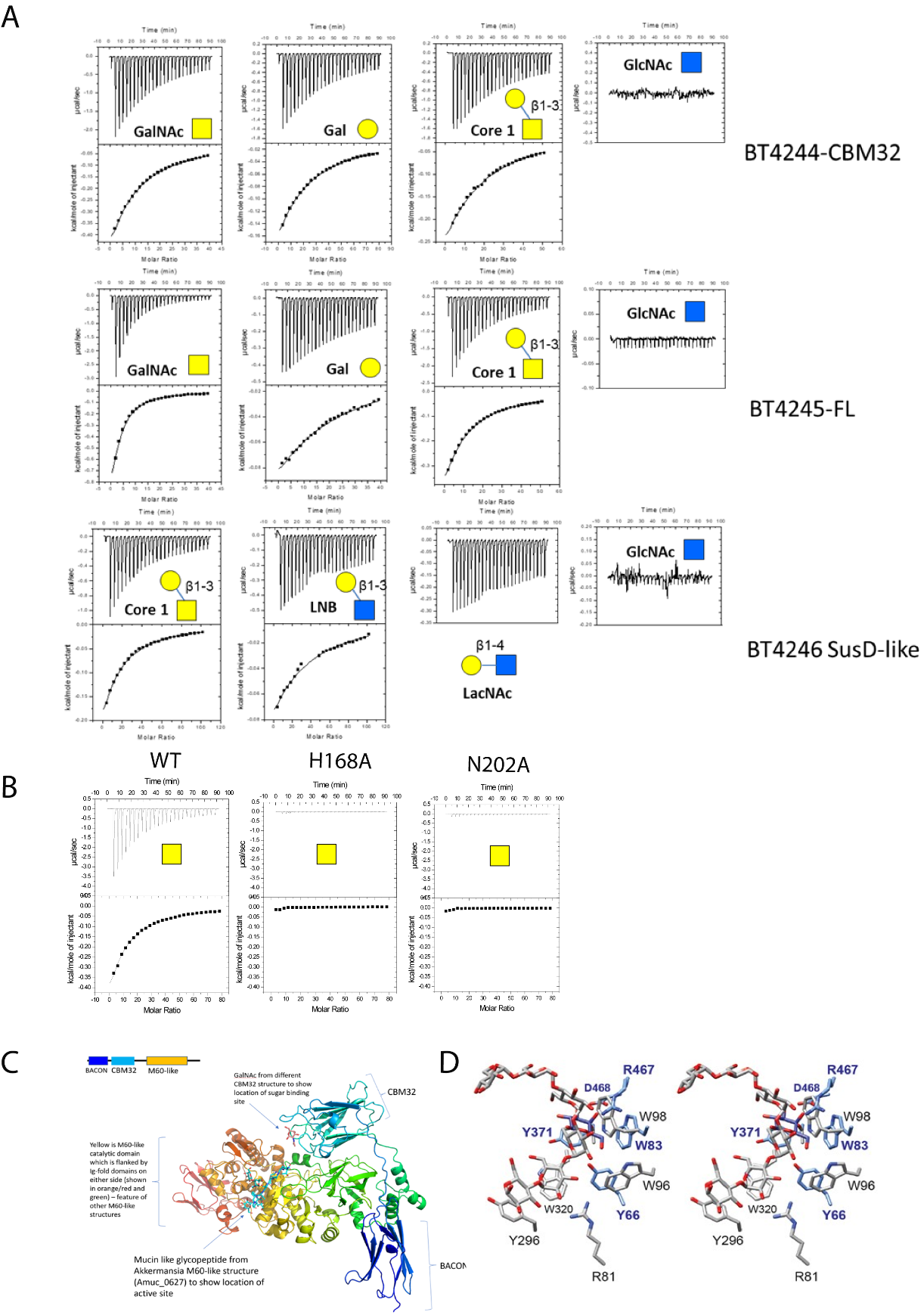

**Supplemental figure 2: Structural and functional insights into mucin sugar recognition by PUL BT4240-50 sugar binding components.** **A:** ITC traces showing binding of PUL encoded BT4244-CBM32 alone, full length SGBP (BT4245) and SusD-like (BT4246) vs mucin derived mono- and di-saccharides. **B:** Effect of BT4244 CBM32 mutations on GalNAc binding **C:** Alphafold2 structure (ColabFold v1.5.5) of full length BT4244-M60L (lacking signal peptide) showing the relative positioning of the various domains (colour coded blue to red N- to C-terminus). The carbohydrate binding site of the CBM32 is oriented such that the CBM could bind to the glycopeptide substrate in the active site of the catalytic domain, potentially contributing to substrate specificity or aiding in substrate positioning in the active site. The glycopeptide shown is overlaid from the structure of an M60-like homologue of BT4244 from *Akkermanisa muciniphila,* Amuc_0627 (PDB 7YX8) **D**: Stereoview superposition of the glycan binding pockets of BT4246 SusD-like and the canonical starch binding SusD, BT3701. Overlay of the Cα backbones of both proteins demonstrate that the ligand-binding site is located in the same position on both proteins, though the chemistry of the site is specific for each glycan. The binding site on BT3701 [82] of the ligand maltoheptaose is shown as grey/red sticks with black residue labels and galactose site of BT4246 is shown as blue/red sticks, with galactose in dark blue, and blue residue labels.

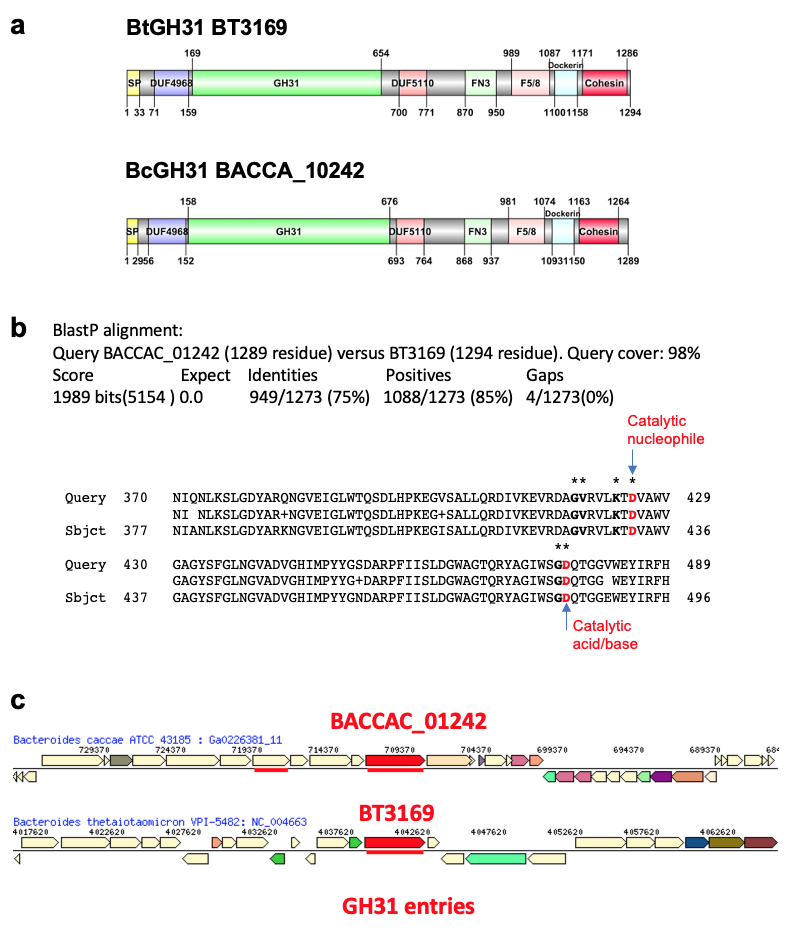

**Supplementary Figure 3. Comparison of the GH31 α-N-acetylgalactosaminidases from *B. caccae* with the most similar homologue from *B. theta*. A:** Comparison of the protein domain organisation of GH31 BACCAC_01242 and BT3169 using DOG2.0 [91] **B:** BlastP alignment between BACCAC_01242 and BT3169 GH31s (75% overall identity between the two proteins) focusing on the segments encompassing the key functional residues as defined for BACCAC_01242 [62]. The two residues in red correspond the two key catalytic residues and the stars indicate additional residues considered important for function [62]. **C:** Comparison of the gene neighbourhood of the two most similar GH31s from B. caccae (BACCAC_01242) and *B. theta* (BT3169) highlighting their distinct, unrelated, genomic configurations. The figure was generated with the “Gene Cart Neighborhoods” tool at the IMG database (<https://img.jgi.doe.gov/cgi-bin/w/main.cgi>).

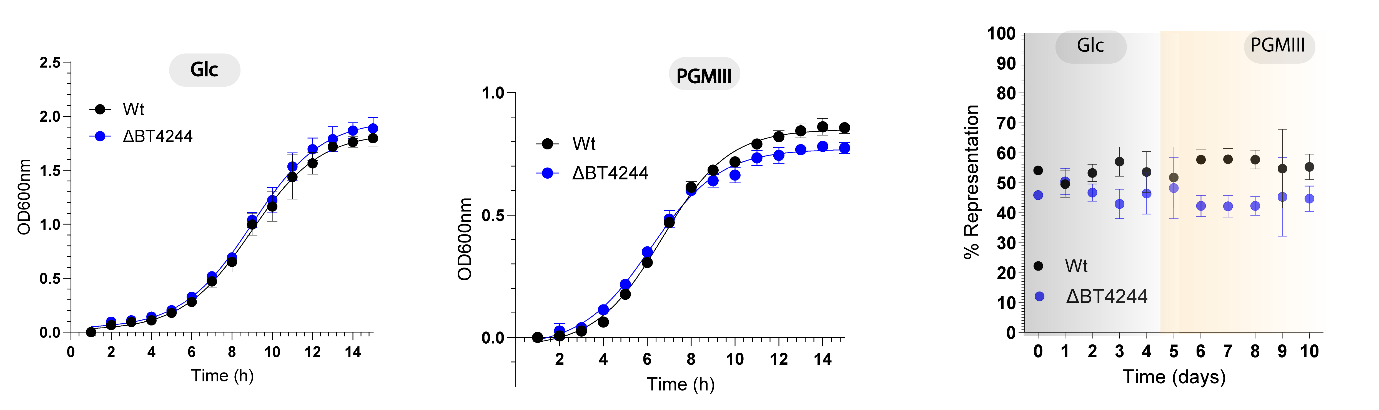

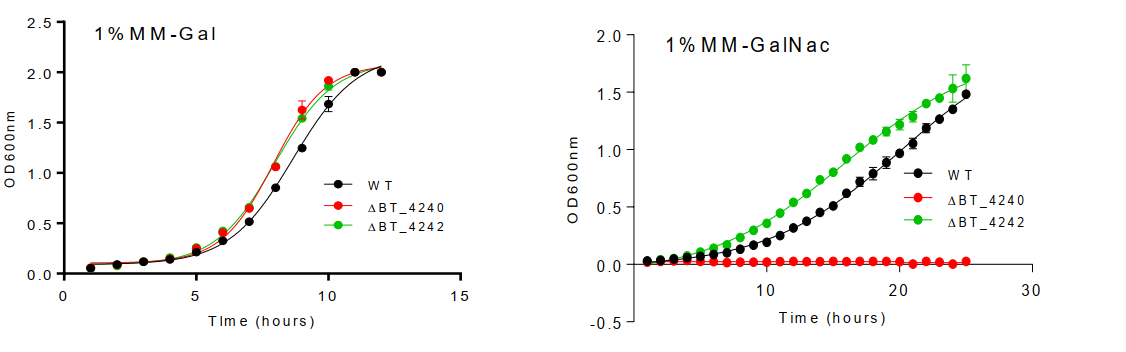

**Supplemental figure 4: Growth profiles of *B.thetaiotaomicron* Wt , ΔBT4244 and ΔBT4242 on glucose and PGMIII and results of competitive growth experiments in Glc and PGMIII.**

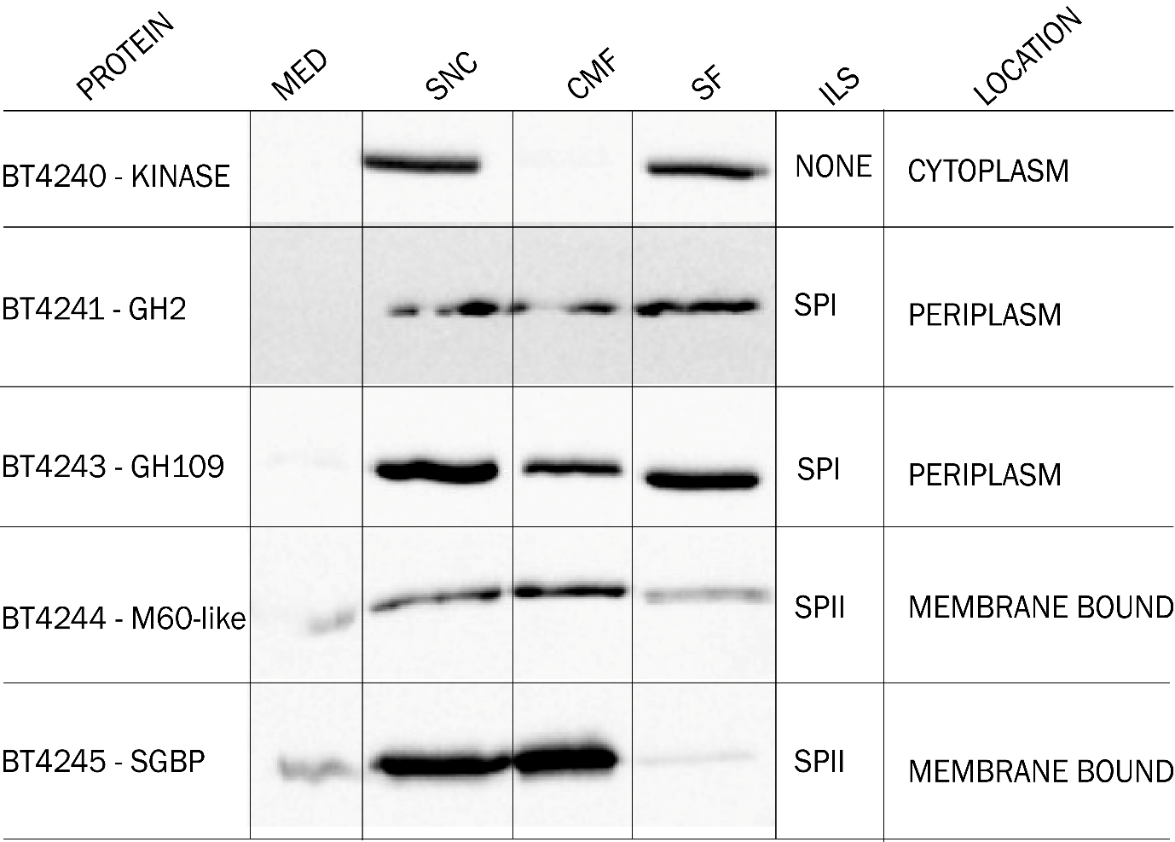

**Supplemental figure 5: Cellular fractionation of PUL BT4240-50 proteins.** Strains of *B. theta* expressing C-terminally Flag-tagged proteins were grown on MM-PGMIII and expression and cellular localization of the tagged proteins analysed by subcellular fractionation and detection using anti-Flag and HRP-conjugated secondary antibodies. MED – spent medium (concentrated to same volume as other fractions), SNC – cell lysate after sonication, CMF – pellet after ultracentrifugation of sonicated cells, SF – supernatant after ultracentrifugation of sonicated cells. ILS – *in-silico* localisation signal based on prediction software LipoP 1.0 (<http://www.cbs.dtu.dk/services/LipoP/>). Location column shows main cellular location based on data shown and predicted signal sequence.

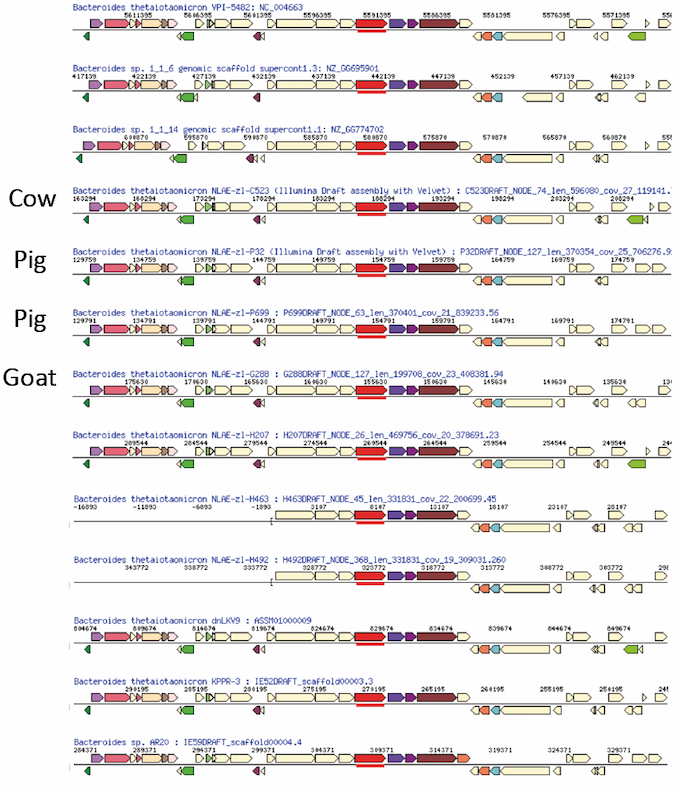

**Supplementary Figure 6. Conservation of PUL BT4240-50 across *B. thetaiotaomicron* isolates from humans and animals.** A total of 14 annotated *B. theta* genomes encoding identical PULs to BT4240-50 were identified using BlastP with BT4244 as query at the IMG database (identity of ≥99% to BT4244). Of these genomes two are derived from the same strain sequence data and only the original annotation is shown (*B. theta* VPI-5482, NC_004663, top entry). The distinct 13 genomes show the same gene set and order (synteny) compared to *B. theta* VPI-5482. For two genomes the scaffolds covering the set of BT4240-50 homologs are partial and do not include the region encoding the three regulatory proteins characteristic of the PUL. All but four strains are from humans with the four animal derived strains being NLAE-z1-C523 (cow), NLAE-z1-P32 (pig), NLAE-z1-P699 (pig), NLAE-z1-G288 (goat). The figure was generated with the “Gene Cart Neighborhoods” tool at the IMG database (<https://img.jgi.doe.gov/cgi-bin/w/main.cgi>).

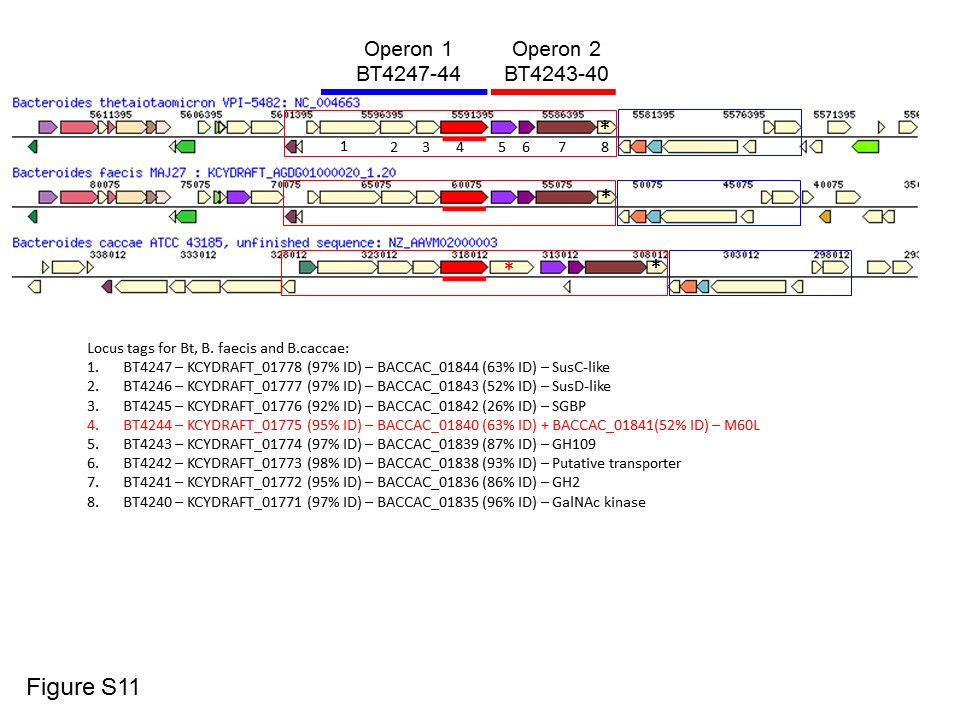

**Supplementary figure 7. Comparison of gene neighbourhood of *B. thetaiotaomicron* BT4240-50 with related PULs from *B. faecis* and *B. caccae*.** In addition to the *B. theta* genomes encoding homologs of PUL BT4240-50, *Bacteroides faecis* MAJ27 and *Bacteroides caccae* ATCC43185 also encode identical (*B. faecis*: KCYDRAFT_01771-81) or very similar, PULs (*B. caccae*: BACCAC_1835-46), with the latter characterised by an additional M60L protease (BT4244 and homologs are shown in red in the three gene neighbourhoods and the additional M60L homolog in *B. caccae* is indicated by the red star). *B. theta* and *B. faecis* share an identical gene neighbourhood organisation across the majority of the shown genome segment. The genes downstream BT4240 and its corresponding homologs from *B. faecis* and *B. caccae* (indicated by a black stars) are conserved between the three species (blue box). The figure was generated with the “Gene Cart Neighborhoods” tool at the IMG database (https://img.jgi.doe.gov/cgi-bin/w/main.cgi). The locus tags, annotations and sequence identity (% values between brackets) between *B. theta* proteins and respectively *B. faecis* and *B. caccae* proteins are listed below the figure.

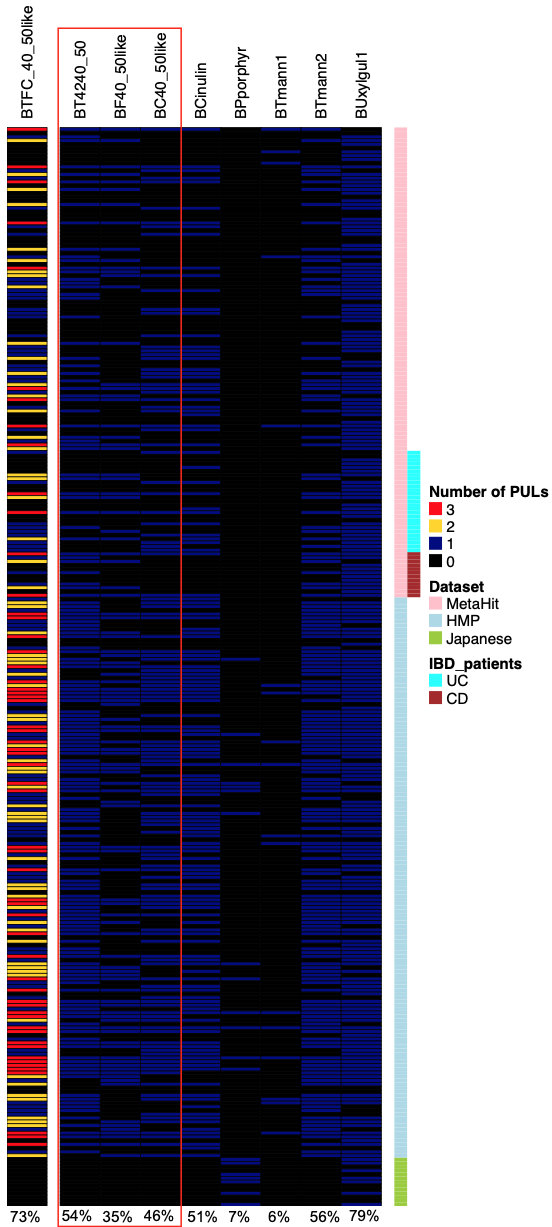

**Supplementary figure 8. Distribution of B. thetaiotaomicron PUL BT4240-50 and the related PUL from *B. faecis* and *B. caccae* across human metagenome datasets.** The human gut metagenome sequence data from 287 individuals were analysed as described previously [24, 25]. The three datasets of human metagenomes were from top to bottom: (i) samples from the MetaHit project [92], (ii) the HMP ([93]) and (iii) 13 healthy Japanese individuals[94] . Samples from inflammatory bowel disease (IBD) patients among the MetaHit dataset are also indicated - ulcerative colitis (UC) and Crohn’s disease (CD). The samples were queried by Blast using DNA sequences from indicated PULs (see methods section for details). Each blue horizontal line represents a positive sample from a given individual. The first column on the left (BTFC40_50like) summarises the sum of the samples positive for the three related BT4240-50 PULs (*B. thetaiotaomicron* BT4240_50), KCYDRAFT_01771-81 (*B. faecies*: BF40_50like) and BACCAC_1835-46 (*B. caccae*: BC40_50like) (See Supplementary fig. 8). The data for these three PULs are highlighted within the red box. The distribution of these three PULs was contrasted with a selection of PULs from various *Bacteroides* species with contrasting frequencies of occurrence across human gut metagenomes: B. caccae inulin PUL (BCinulin, common) [27], *B. plebeius* porphyran PUL (BPporphyr, rare outside Japanese) [95]; *B. theta* mannan 1 PUL (BTmann1, rare) and *B. theta* mannan 2 PUL (BTmann2, common) [24]; and *B. uniformis* xyloglycan PUL (BUxylglul1, very common) [25]. The frequency of a given PUL across the 287 samples is shown at the bottom of each column.
